# TCRpcDist: Estimating TCR physico-chemical similarity to analyze repertoires and predict specificities

**DOI:** 10.1101/2023.06.15.545077

**Authors:** Marta A. S. Perez, Johanna Chiffelle, Sara Bobisse, Francesca Mayol-Rullan, Marion Arnaud, Christophe Sauvage, George Coukos, Alexandre Harari, Vincent Zoete

## Abstract

Approaches to analyse and cluster TCR repertoires to reflect antigen specificity are critical for the diagnosis and prognosis of immune-related diseases and the development of personalized therapies. Sequence-based approaches showed success but remain restrictive, especially when the amount of experimental data used for the training is scarce. Structure-based approaches which represent powerful alternatives, notably to optimize TCRs affinity towards specific epitopes, show limitations for large scale predictions. To handle these challenges, we present TCRpcDist, a 3D-based approach that calculates similarities between TCRs using a metric related to the physico-chemical properties of the loop residues predicted to interact with the epitope. By exploiting private and public datasets and comparing TCRpcDist with competing approaches, we demonstrate that TCRpcDist can accurately identify groups of TCRs that are likely to bind the same or similar epitopes. Additionally, we experimentally validated the ability of TCRpcDist to predict antigen-specificities of tumor-infiltrating lymphocytes orphan TCRs obtained from four cancer patients. TCRpcDist is a promising approach to support TCR repertoire analysis and cancer immunotherapies.

**One Sentence Summary:** We present a new approach for TCR clustering which allows TCR deorphanization for the first time.

## Introduction

For future advancements in cancer immunotherapy, promising approaches such as adoptive cell therapy or vaccination would require the precise knowledge of which T-cell receptor (TCR) recognizes which cancer epitope (*i.e.* pMHC) in a given patient’s tumor(*1*). Although the principle of the binding of TCRs to pMHC has been established long time ago(*2, 3*), it remains extremely challenging to predict which TCRs are responsible for an antigen response. There are several reasons for this. First, the process of V(D)J recombination has the potential to generate an astronomical number of distinct TCRs with biological significance, estimated to be up to 10^20^ (*4*). Furthermore, the binding between TCRs and cancer-related pMHC is of relatively low-affinity(*5, 6*) and is degenerate, meaning that several different TCRs can recognize the same pMHC while at the same time one given TCR can recognize distinct pMHCs. Despite recent sequence- and structure-based computational advances(*7, 8*), the unambiguous prediction of TCR-pMHC pairing, from pools of thousands of candidates, remains a daunting task(*9*).

Recent computational studies demonstrated that common patterns can be identified among TCR sequences interacting with the same epitope(*10–18*) opening the road to an in silico characterization of the specificity, diversity and complexity of TCR repertoires obtained experimentally from one or several patients(*4, 19*). Such approaches are already widely employed (*20–23*).

Analysis of TCR sequences using a panel of pMHC-multimer sorted cells and TCR-pMHC 3D structures led Glanville et al. in 2017 (*10*) to the conclusion that determining possible pMHC contact sites in CDR3s, notably in CDR3β, would offer an opportunity to cluster TCRs with a high probability of sharing the same specificity. Based on this assumption, the authors developed the GLIPH program (grouping of lymphocyte interactions by paratope hotspots) to cluster TCRs based on global and local TCR sequence similarity. Clusters can be filtered for shared V-gene usage, resulting in GLIPH specificity groups, likely to recognize the same or very similar pMHC ligands. The latter are scored based on the enrichment of common V-genes, the CDR3 lengths, clonal expansions, shared HLA alleles among contributors, motif significance and cluster size. When benchmarking GLIPH on a training set of 2,068 unique sequences spanning eight pMHC specificities, the authors found that when combining local and global similarity 14% of the TCRs were clustered, of which 94% were correctly grouped with other TCRs of common specificity. Such an approach could be used to cluster TCRs that recognize the same epitope and to predict their HLA restriction. However, GLIPH loses efficiency and accuracy when more than 10’000 TCRs are analyzed. At this size, artifactual clusters caused by “small-world” effect begin to form. (*24*). To circumvent this problem Huang et al. developed GLIPH2(*24*) in 2020. The latter can process millions of TCR sequences. Among other improvements, in GLIPH2, the “small world” effect was solved by restricting TCR members to be of the same length for a cluster based on global similarity and to differ at the same position between exchangeable amino acids according to a BLOSUM62 matrix. For a cluster based on local similarity, they restricted the CDR3β position to only vary within three amino acids.

Dash et al. in 2017(*11*) defined a distance measure on the space of TCRs, TCRdist, allowing clustering and visualization of repertoire diversity. This quantitative measure of similarity between paired αβ TCRs is obtained by listing the residues belonging to the CDR1, 2 and 3 loops, as well as an additional variable loop between CDR2 and CDR3 since they are known to possibly contact the pMHC, and by computing a similarity-weighted mismatch distance defined based on the BLOSUM62 substitution matrix, with a gap penalty to capture variations in the length of the CDRs. Of note, a higher weight was given to the CDR3 sequence in view of its prominent role in epitope binding. This distance can then be calculated for each possible pair of TCRs belonging to a given repertoire, generating a so-called distance matrix. The latter can be used for TCRs clustering, or the construction of hierarchical distance trees to analyze the diversity and complexity of the TCR repertoire. In 2021, Mayer-Blackwell et al. (*22*) used a new version of TCRdist, TCRdist3(*25*), to guide the formation of meta-clonotypes (i.e. groups of TCRs with biochemically similar CDRs that likely share antigen recognition) optimized for biomarker development. TCRdist3 brings new flexibility to distance-based repertoire analysis, allowing customization of the distance metric, analysis of 8ψ TCRs, and at-scale computation with sparse data representations and parallelized calculations.

In 2019, Ostmeyer and colleagues (*12*) introduced an approach that consists in feeding machine learning techniques, based on logistic regressions, with biophysicochemical descriptors of the TCR interface for analyzing immune repertoires of several patients, with the objective to identify differences between TCRs in normal and tumor tissues. In this approach, the biophysicochemical characteristics of sliding windows of 4 consecutive residues of CDR3 β (i.e. the so called 4-mers), excluding the first 4 and last 3 residues, are described using five Atchley factors(*26*) that encode for codon diversity, secondary structure, molecular size, polarity, and electrostatic charge of the residues. The method identified a short list of preferred values for these descriptors at key positions in TCRs present in the tumor, which permitted the identification of disease-associated TCRs. Although this approach must be retrained for each set of TCRs studied and is restricted to CDR3β only (which limits its predictive ability), this type of sequence-based ‘property’-based approach could circumvent some of the drawbacks of purely sequence-based analysis, such as the need to have very large numbers of disease-associated TCR sequences available for training, and the possibility of detecting potential antigen-binding TCRs even if their sequence differ from those that have been previously encountered.

Also in 2019, Lanzarotti et al developed a model for prediction of TCR targets based on similarity to a database of TCRs with known pMHC (*27*). They showed increased predictive ability by focusing on CDRs rather than the full length TCR protein sequences, by incorporating information from paired α and β chains, and by integrating information for all 6 CDR loops rather than just CDR3. Additionally, they demonstrated that the inclusion of structural information in the model improved, consistently yet modestly, the accuracy of the epitope prediction, in particular in situations where no sequence with high similarity is available in the TCR database. They expected this to evolve favorably, as the accuracy of TCR structural modeling tools improve (*28–30*) and the number of available TCR 3D structures to be used as modelling templates increases (*31*).

In 2021, Ehrlich introduced SwarmTCR, a method that combines sequence-based approach with structural information (*32*). SwarmTCR uses CDRs sequences to predict the specificity of TCR using a nearest-neighbor approach. The approach works by optimizing the weights of the individual CDR regions to maximize classification performance, a knowledge taken from TCR:pMHC structures (*33*). CDRs are thus weighted differently, depending upon the peptide being presented and the MHC type, to better describe the involvement of TCR in binding to the pMHC. SwarmTCR showed comparable performance to TCRdist when using TCR sequence information from single cell and bulk data recognizing 5 different pMHCs (*32*). The performance, robustness and generalizability of this approach is highly dependent on the training data and, therefore, it will be difficult to extend its use to pMHCs without or with a very limited number of known binding TCRs.

Approaches predicting TCR-pMHC binding based on their 3D structure and on force-field based modelling have already been investigated (*34–36*). Although physics-based Molecular Modelling approaches at the atom scale have been proven powerful to optimize the affinity of a TCR towards a specific epitope, they are limited by calculation speeds and therefore can only be applied to a limited number of TCRs. Very recently Lin et al. introduced RACER (Rapid Coarse-Grained Epitope TCR), a pairwise energy model capable of rapidly assessing TCR-peptide affinity for large scale repertoires, the MHC being constant (*36*). As a Coarse-Grained model each residue is not fully described at the atom level but instead defined by the positions of its three atoms Cα Cβ and O (except for GLY that do not have a Cβ). RACER applies supervised machine learning to distinguish strong TCR:pMHC pairs from weak pairs, with fixed MHC. The trained parameters further enable a physical interpretation of the interaction patterns encoded in TCR. When structural data for a specific TCR:pMHC pair is unavailable, they built the TCR:pMHC models based on TCR:pMHC experimental structures (MHC is kept fixed). Structural relaxation data is needed for structural models because TCR:peptide clashes disfavor strong binders and poses a challenge for large scale analysis. Despite the fact that this approach depends on the training data and that pairwise interactions are only one of several factors influencing epitope recognition, this tool can be used to understand general questions regarding TCR and relevant antigen landscape. It would be, however, difficult to use this approach for TCR and pMHC pairing from a pool of thousands of candidates, since (i) this would require making a model for all possible TCR:pMHC combinations including structural relaxation, and because (ii) the scope would be limited by the fact that the approach requires treating the different alleles separately and the TCR:pMHC structure of the allele of interest has to exist on Protein Data Bank (PDB)(*33, 37*).

By improving from the above approaches and circumventing their limitations, we developed TCRpcDist, a novel and rapid approach that allows TCR clustering by analyzing the biophysicochemical properties of the solvent exposed 4-mer residue motifs of CDR1α, CDR2α, CDR3α, CDR1β, CDR2β, and CDR3β. TCRpcDist was compared with other existing approaches and further validated using public and private TCR data sets. TCRpcDist can contribute identifying: (1) clusters of TCRs that are likely to bind the same or similar known epitopes; (2) the most probable epitope targeted by an orphan TCR among a pool of pMHCs for which TCR binders are known.

## RESULTS

### TCRpcDist using only CDR3β loops permits TCRs clustering and specificity correlation

It is generally admitted that the CDR3β loop of a TCR makes most of the interactions with the peptide of a cognate pMHC(*38, 39*). Accordingly, many TCR specificity predictors focus only on the CDR3β sequence. Therefore, our analysis was first carried out by making the same assumption and clustering the TCRs using an Atchley-based distance calculated on sliding windows of 4 consecutive residues (i.e. the 4-mers) extracted only from the CDR3β of each TCR. To avoid limitations that could be generated by uncertainties in 3D-structure models (when calculating nSESA, see below), the approach was initially tested using 54 TCRs taken from experimentally determined structures of TCR:pMHC complexes stored in the PDB (**SI Table 1**). These TCRs recognize 16 different known pMHCs. The calculated clustering is depicted in **FIG. 1a**. Note that the clustering shown in **FIG. 1a** was obtained according to the Atchley-based distance between TCRs CDR3β only, without any information regarding cognate pMHCs. The coloring according to the TCR specificity is only applied afterwards, to display if similar TCRs indeed bind identical pMHC. These initial results show that, even though only CDR3β residues are used to calculate distances, clusters of receptors binding the same pMHC tend to spontaneously form.

**FIG. 1.**
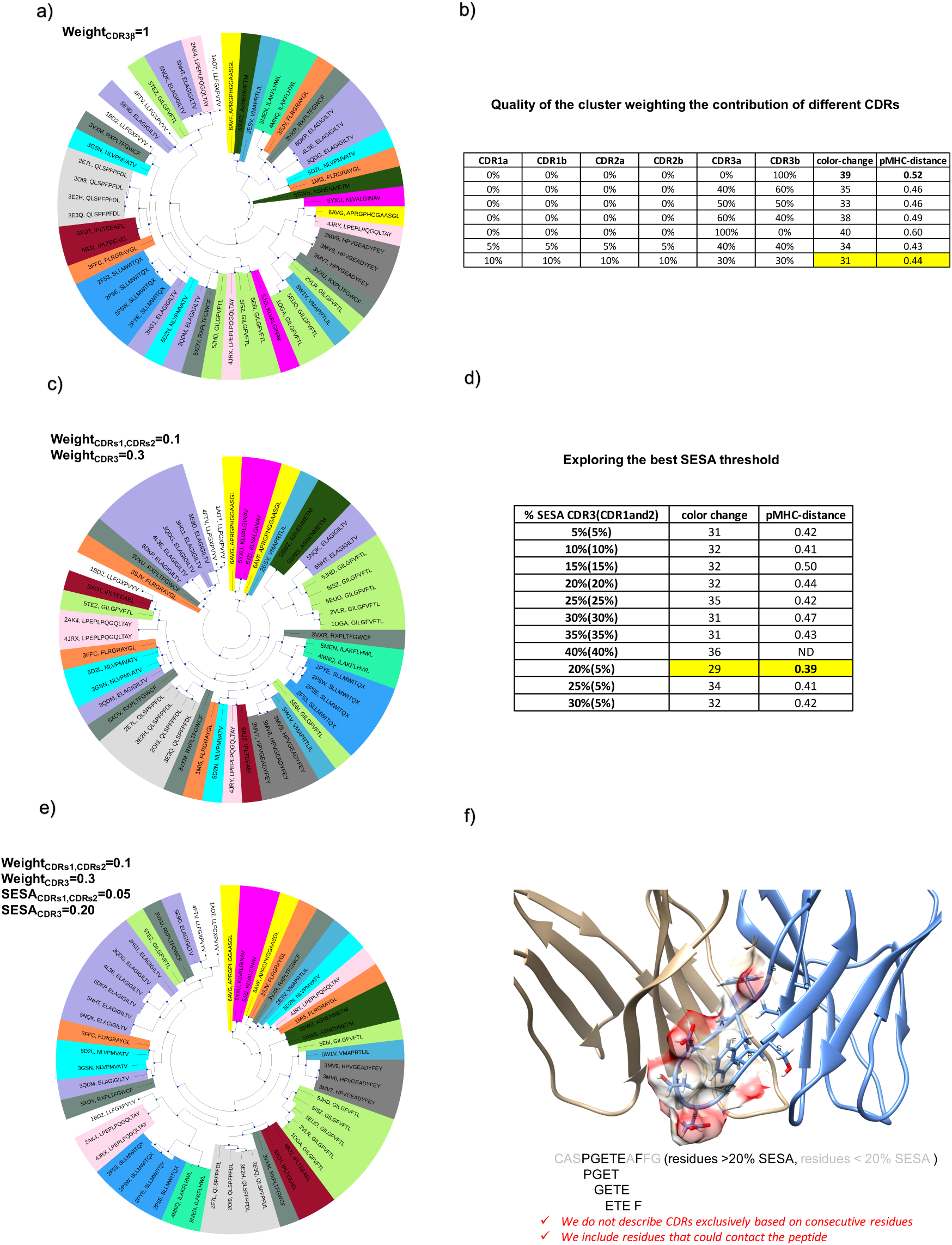
TCRpcDist clustering TCRs and correlating with their specificity. a) shows hierarchical clustering of a set of 54 TCRs recognizing 16 different pMHC using the Atchley-based distance considering only sliding windows of 4 consecutive residues of the CDR3β. After clustering, each TCR is colored according to the pMHC it binds. The sequence of the bound peptide is also given; b) quality of the cluster as measured by the number of color changes and the pMHC-distance, for diverse weightings of the contributions of the various CDRs. The maximal clustering efficiency is highlighted in yellow and obtained when each CDR3s contribute by 30% and each of the remaining CDRs by 10% to the distance calculation; c) shows the hierarchical clustering of a set of 54 TCRs recognizing 16 different pMHC using the Atchley-based distance considering all 6 TCR CDRs (*i.e.*, CDR1α, CDR2α, CDR3α, CDR1β, CDR2β, and CDR3β). After clustering, each TCR is colored according to the pMHC it binds. The sequence of the bound peptide is also given; d) exploring the best nSESA threshold. The clustering efficiency as measured by the number of color changes and the pMHC-distance is maximal when residues with nSESA < 5% in CDRs 1 and 2 and residues with nSESA < 20% in CDRs 3 are excluded from the distance calculation; e) shows hierarchical clustering of a set of 54 TCRs recognizing 16 different pMHC using the Atchley-based distance considering all 6 TCR CDRs (*i.e.*, CDR1α, CDR2α, CDR3α, CDR1β, CDR2β, and CDR3β), as well as residues buriedness. After clustering, each TCR is colored according to the pMHC it binds. The sequence of the bound peptide is also given; f) illustrates how solvent exposed residues are included in the distance calculation while buried residues are excluded (The TCR structure corresponds to PDB ID 4JRX).

We measured the number of times the color changed between two successive nodes in the hierarchical tree (**FIG. 1a**) as a measure of the clustering quality, starting from the upper node and turning clockwise. 39 color changes are observed, which is an encouraging result in view of the fact that a random clustering provides an average of 50.7 color changes (p<0.0001). The quality of the clustering was also determined by a more quantitative metric, pMHC-distance, defined as the average branch length distance between all possible pairs of TCR nodes that recognize the same pMHC. The lower this average, the closer the TCRs binding the same pMHC are grouping in this hierarchical clustering. The pMHC-distance value is 0.52 for the tree generated by TCRpcDist and CDR3β loops only, while a random clustering provides a pMHC-distance of 0.94 (p<0.0001; See **Table 1)**. Of note, each pMHC contributes with the same weight (1/16) to the final pMHC-distance, to prevent this measure to be biased by the most frequent pMHCs.

**Table 1.**
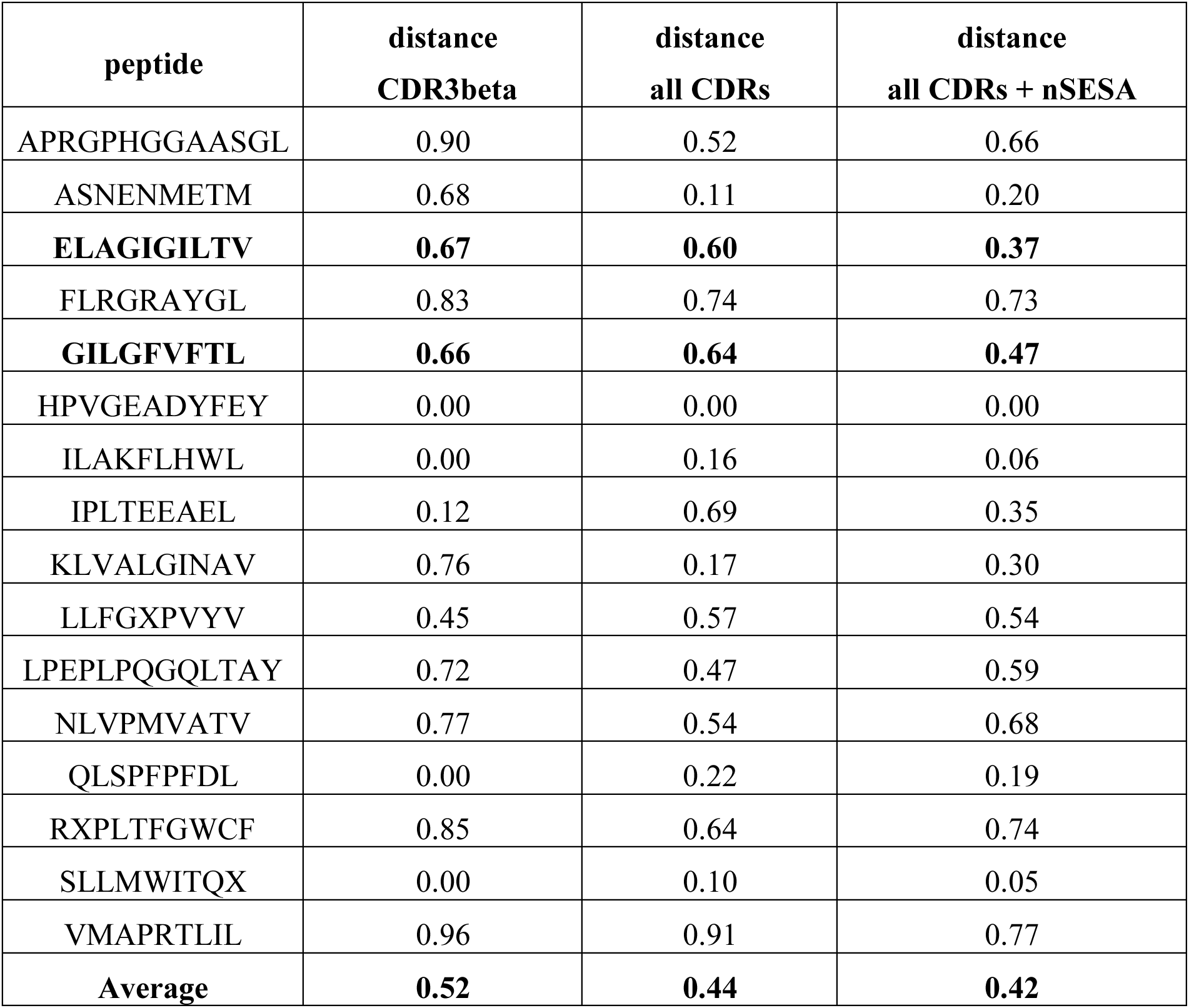
Average branch length distance per peptide in the hierarchical tree for the PDB set considering three different scenarios: considering only CDR3β, or all CDRs, or all CDRs+ nSESA in the distance calculation. Each peptide contributes with the same weight (1/16) to the final pMHC-distance, to prevent this measure to be biased by the most frequent pMHCs.

| peptide | distance<br>CDR3beta | distance<br>all CDRs | distance<br>all CDRs + nSESA |
| --- | --- | --- | --- |
| APRGPHGGAASGL | 0.90 | 0.52 | 0.66 |
| ASNENMETM | 0.68 | 0.11 | 0.20 |
| **ELAGIGILTV** | **0.67** | **0.60** | **0.37** |
| FLRGRAYGL | 0.83 | 0.74 | 0.73 |
| **GILGFVFTL** | **0.66** | **0.64** | **0.47** |
| HPVGEADYFEY | 0.00 | 0.00 | 0.00 |
| ILAKFLHWL | 0.00 | 0.16 | 0.06 |
| IPLTEEAEL | 0.12 | 0.69 | 0.35 |
| KLVALGINAV | 0.76 | 0.17 | 0.30 |
| LLFGXPVYV | 0.45 | 0.57 | 0.54 |
| LPEPLPQGQLTAY | 0.72 | 0.47 | 0.59 |
| NLVPMVATV | 0.77 | 0.54 | 0.68 |
| QLSPFPFDL | 0.00 | 0.22 | 0.19 |
| RXPLTFGWCF | 0.85 | 0.64 | 0.74 |
| SLLMWITQX | 0.00 | 0.10 | 0.05 |
| VMAPRTLIL | 0.96 | 0.91 | 0.77 |
| **Average** | **0.52** | **0.44** | **0.42** |

### TCRpcDist is more efficient when using all CDRs (α and β chains) compared to CDR3β loops only

The claim that CDR3β is the most important contributor to TCR specificity is a limiting approximation, as several approaches have already pointed out (*11, 32*). Indeed, we could evaluate the relevance of the other CDRs residues in the peptide recognition in Figure 1 of a previous study (*7*). Consequently, we decided to consider the contributions of all six loops, *i.e.*, CDR1α, CDR2α, CDR3α, CDR1β, CDR2β, and CDR3β in our TCR distance definition. The quality of the resulting clustering as measured by the color change and the pMHC-distance in the hierarchical tree was analyzed by weighting the contribution of the different CDRs (**FIG. 1b)**. The clustering relevance is maximal when weighting values of 30% are applied to the subset of amino acids in CDR3α or CDR3β and of 10% to the subset of amino acids in CDR1α, CDR2α, CDR1β, or CDR2β (See **FIG. 1b**). To avoid overfitting the model on the training set, we did not perform a full sampling of the effects of the weights to optimize their values. **FIG. 1c** shows the corresponding hierarchical clustering. Of note, this clustering is much better at grouping together TCRs that bind the same pMHC, with only 31 color changes (p<10^-6^ compared to random), compared to 39 when considering only CDR3β (**FIG. 1a**). Concomitantly, the pMHC-distance decreases to 0.44, compared to 0.52 when considering only CDR3β (**Table 1**). For example, the 3 TCRs recognizing the peptide LPEPLPQGQLTAY, with PDB codes 4JRY, 4JRX and 2AK4 are not clustered when we perform the analysis based exclusively on CDR3n (pink slices are separated in **FIG. 1a**). The clustering efficiency improves when considering all the CDRs, with 2 TCRs (2AK4 and 4JRX) out of 3, now grouped together (**FIG. 1b**). This results from the fact that they share the same TRAV, CDR3α and TRBV genes, i.e., same CDR1α, CDR2α, CDR3α and CDR1β and CDR2β (**SI Table 1**, TCRs with PDBID 2AK4 and 4JRX). Another illustration of the improved efficiency comes from the TCR with PDB ID 1A07 that recognizes the peptide LLFGXPVYV but which wrongly clusters with 2AK4, a TCR recognizing the peptide LPEPLPQGQLTAY, when using uniquely the CDR3β (pink and white slices next to each other in **FIG. 1a**). The misleading cluster when using exclusively CDR3n can be explained by the fact that they share the 4-mer feature GLAG in CDR3β. 1A07 and 2AK4 cluster properly when considering all the CDRs (See **FIG. 1b**), again due to the importance of the other CDRs on top of the CDR3β in pMHC recognition and therefore in clustering.

We note that when working with bulk sequencing data, we do not have αβ pairing information and, despite the lower accuracy of the approach using uniquely CDR3β information, we could still successfully use it for clustering purposes, with results much better than random (p-value <0.0001). The relevance of the clustering approach using exclusively CDR3β residues was already shown in a previous study (*7*).

### The maximal clustering efficiency of TCRpcDist-3D is obtained when taking the solvent accessibility of the CDRs residues into account

Another route of enhancement is to take the solvent accessibility of the CDRs residues into account. Indeed, even though they belong to CDRs, residues that are buried into the TCR structure are not available for interaction with the pMHC. Thus, it is possible to enhance the clustering approach by removing the buried residues from those considered in the distance calculation.

Therefore, normalized Solvent Excluded Surface Area (nSESA) for all residues in the 6 CDRs were calculated for the experimental 3D structures of the above-mentioned 54 TCRs retrieved from the PDB. The clustering efficiency was tested on different SESA thresholds (**FIG. 1d**). Then, the residues of CDRs 1 and 2 (α and β), with a nSESA > 5% were considered for distance calculation, while a threshold of 20% was applied for the two CDR3s (α and β), the combination of parameters that give the maximum clustering efficiency. Accordingly, the distances between CDRs were calculated using sliding windows of 4 consecutive solvent-exposed residues (and not just consecutive residues) for this test set of 54 TCRs. Taking the buriedness of the residues into account enhanced the quality of the clustering (**FIG. 1e**). The number of color changes in the clustering tree reached 29 and the pMHC-distance decreased to 0.42 (p<0.0001 compared to random and compared with previous versions of the approach) (see **Table 1**).

Noticeably, we can see in **FIG. 1f** that the 4^rd^ residue of the CDR3β, in this example, i.e. Pro, excluded from the distance calculation in the previous version of the approach, is sufficiently solvent exposed and should be considered for the analysis of the TCR distance. Indeed, the 3D structure of the complex TCR:pMHC PDB ID 4JRX shows that the Pro residue contacts the peptide (**SI Figure 1**). This illustrates that approaches which systematically get rid of the first 4 and the last 3 residues from the CDR loops (*12, 23*), since they are supposed to be likely buried, might in fact accidentally remove residues relevant for binding. Our approach tries to avoid this approximation by computing the nSESA for all CDR residues to determine their solvent accessibility and excluding the buried residues (nSESA < 20%) from the distance calculation.

Analyzing the clustering trees (**FIG 1a, 1c, 1e**) we observe that, for example, the TCRs recognizing the peptide ELAGIGILTV cluster much better when solvent accessibility is considered (purple slices closer in **FIG. 1e**). The average of the branch length distance between the nodes of ELAGIGILTV-binding TCRs decreases from 0.67 (using just CDR3β) to 0.60 (using all CDRs) and ultimately to 0.37 (when using nSESA on top of all CDRs, see **Table 1)**.

### Clustering efficiency of TCRpcDist-3D using a private data set of TCRs

The approach was then applied to a private set of 45 TCRs with 12 different known cognate pMHCs (some described in Arnaud et al. (*40*), **SI Table 2**, see methods). Since the experimental 3D structure of these TCRs is unknown, the buriedness of all the CDR residues was calculated based on TCR models. The clustering trees for this assessment set are depicted in **FIG. 2** and as for the previous set, we measured the quality of the cluster by color change and pMHC-distance. The number of color changes was 33 using only CDR3β (**FIG. 2a)**, 33 considering contribution of all CDRs (**FIG. 2b)** and 27 considering all CDRs and nSESA (**FIG. 2c),** with results much better than random (p-value <0.0001). The pMHC-distance was 0.33, 0.31 and 0.30 for these three levels of approximation, respectively. Once again, we observe an increase of the clustering efficiency using all the CDRs and nSESA (TCRpcDist-3D), confirming the robustness of the approach, even though we are using TCR structural models instead of TCR experimental structures. Calculation of TCR loop conformations being stochastic, TCRpcDist-3D is itself stochastic, contrarily to TCRpcDist which is deterministic. However, the variability on calculated TCRpcDist-3D distances between several runs, starting from the same input, remains very limited (See **SI VariabilityOfOutcome.xlsx**) and does not change the conclusions of TCR distance analysis.

**FIG. 2.**
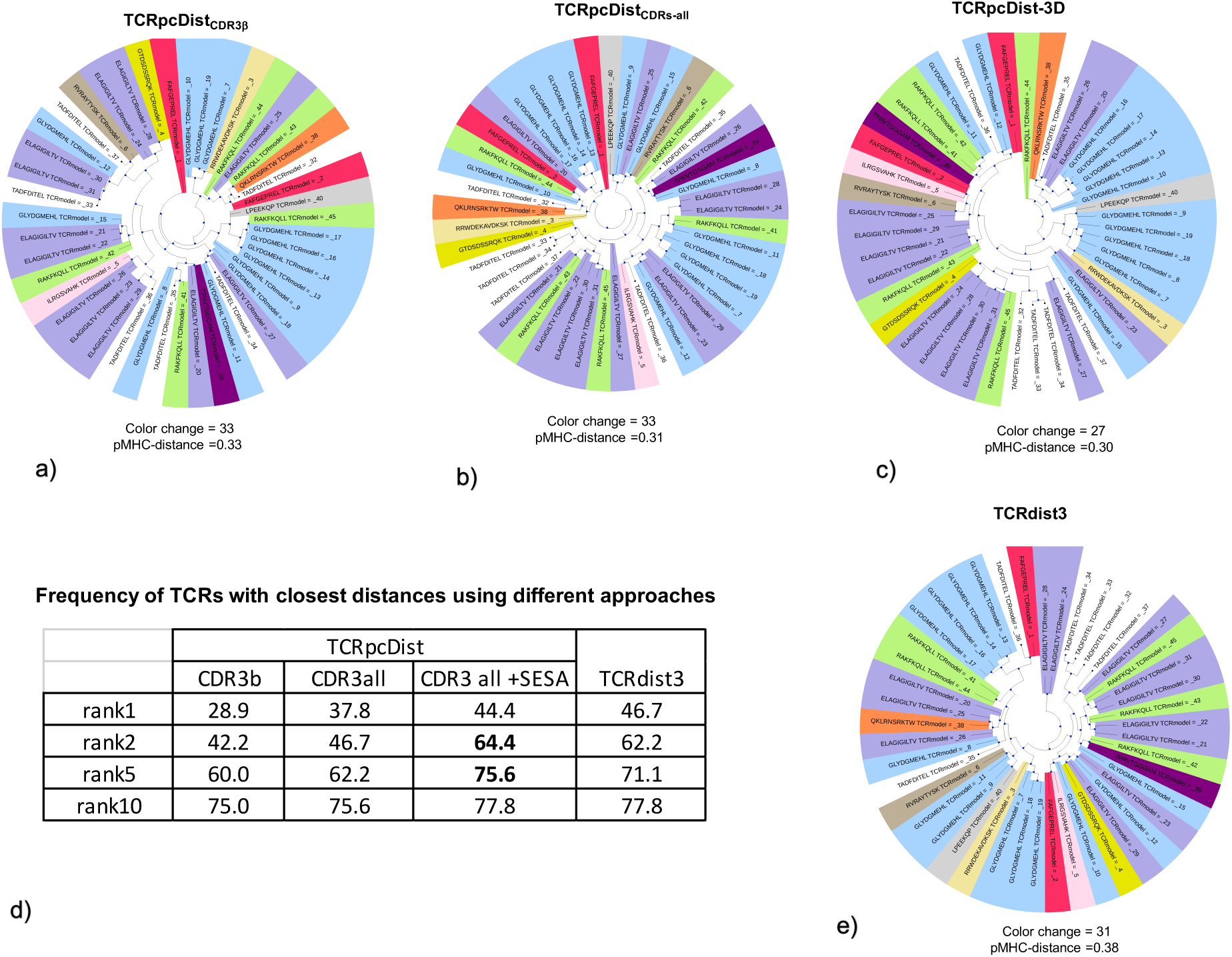
TCRpcDist clustering TCRs and correlating with their specificity using a private set of TCRs with known specificity a) shows the hierarchical clustering of a test set of 45 TCRs recognizing 12 different pMHC using the Atchley-based distance considering only sliding windows of 4 consecutive residues of the CDR3β. These TCRs were not used to choose the TCRpcDist parameters. After clustering, each TCR is colored according to the pMHC it binds. The sequence of the bound peptide is also given. b) shows the hierarchical clustering of a set of 45 TCRs recognizing 12 different pMHC using the Atchley-based distance and considering all 6 TCR CDRs (*i.e.*, CDR1α, CDR2α, CDR3α, CDR1β, CDR2β, and CDR3β). After clustering, each TCR is colored according to the pMHC it binds. The sequence of the bound peptide is also given. c) shows the hierarchical clustering of a set of 45 TCRs recognizing 12 different pMHC using the Atchley-based distance, considering all 6 TCR CDRs (*i.e.*, CDR1α, CDR2α, CDR3α, CDR1β, CDR2β, and CDR3β) as well as residues buriedness. After clustering, each TCR is colored according to the pMHC it binds. The sequence of the bound peptide is also given d) Table shows how often a TCR with the same specificity is found in the top 1, 2, 5 and 10 TCRs with the closest distances using the 3 versions of TCRpcDist and using the distances from the TCRdist3 approach e) shows hierarchical clustering of a set of 45 TCRs recognizing 12 different pMHC using TCRdist3 (*22*)

Analyzing the clustering trees (**FIG. 2a, 2b, 2c**) we observed that, for example, the TCRs recognizing the peptide ELAGIGILTV (overrepresented in the set with 27% of the TCRs, **SI Table 2**) cluster much better with TCRpcDist-3D (purple slices closer in **FIG. 2c**). Nevertheless, even at this level of approximation (**FIG. 2c**) they tend to form sub-clusters instead of a single cluster. On the contrary, the TCRs recognizing the same peptide - also overrepresented in the PDB (17% of the TCRs, **SI Table 1**) - form mainly a single cluster when applying TCRpcDist-3D on the PDB set (**FIG. 1e**). The fact they form sub-clusters in the present set is justified by a much higher TCRs sequence variability. TCRs recognizing ELAGIGILTV in this set exhibit 10 different TRAV genes, 8 different TRBV genes, 18 different AA in CDR3α and 16 in CDR3β, while TCRs recognizing ELAGIGILTV in the PDB set show only 2 different TRAV genes (87.5 % of the TCRs recognizing TRAV12-2), 5 different TRBV genes and 12 different AA in CDR3α and β. The statistics regarding the different composition of the TCRs recognizing ELAGIGILTV are given in **SI Table 4**. The fact that the present set contains singleton TCRs and much higher sequence variability for TCRs recognizing the same peptide makes the clustering more challenging. Still the clustering efficiency is remarkable here and specially for TCRpcDist-3D. Comparison with other methods confirm the high accuracy of our approach (see section the next section “TCRpcDist-3D compares favorably to state-of-the-art approaches”).

Additionally, we have studied how frequently, starting from a given TCR, it is possible to find another TCR with the same specificity in the 1, 2, 5 and 10 top-ranked TCRs with closest distances. TCR pairs with the closest distance (rank 1) share the same specificity in 28.9% of the cases using only CDR3β, 37.8% considering contribution of all the CDRs and in 44.4% of the cases considering all CDRs and solvent accessibility. In all cases, these success rates are much better than what can be obtained by random assignments (19.0% success; p-value<0.0001). The predictive power of our approach is increased if we consider the top 2, top 5 or the top 10 closest TCRs to a given TCR (**FIG. 2d**). Indeed, the specificity of a given TCR is shared by at least one of the 2, 5 and 10 top-ranked TCRs in 42.2%, 60.0% and 75% of the cases, respectively, when using exclusively CDR3β (p-value < 0.0001). When considering all the CDRs, the success increased to 46.7%, 62.2% and 75.6%, and even reached 64.4%, 75.6% and 77.8% considering all CDRs and nSESA. The gain is impressive when comparing the data with and without nSESA (maximum gain of 22.2% points for rank2 when comparing CDR3β with all CDRs and nSESA). We note that this set contains singleton peptides for 6 TCRs, for which is therefore impossible to find a TCR pair with the same specificity, hence the maximum probability of finding a TCR pair is 86.7% and not 100%. As for this study we have used structural models instead of experimental structures from the PDB, we thought this could limit the efficiency of the 3D/nSESA addition. However, once again, the superior accuracy of the version of the approach that includes all CDRs and nSESA (TCRpcDist-3D) was confirmed.

### TCRpcDist-3D compares favorably to state-of-the-art approaches

We applied one of the most renown and freely available method, TCRdist3, to analyze the diversity and complexity of the private 45-TCRs repertoire and compare with TCRpcDist-3D, **FIG 2e**. We chose this repertoire since its content was not used to train either of these two approaches, ensuring an unbiased comparison. When both approaches are applied to this private set of 45 TCRs with known specificities to cluster them in the form of a hierarchical tree, TCRdist3 yields 31 color changes and a pMHC-distance of 0.38, while TCRpcDist-3D yields 27 color changes and a pMHC-distance of 0.33. This significant difference in the number of color changes and pMHC-distance values (p<0.0001), indicates a slightly higher clustering efficiency for the TCRpcDist-3D based approach. To evaluate if we can use both TCRpcDist-3D and TCRdist3 in a consensual way, and possibly generate a synergy between the two approaches, we have combined TCRpcDist-3D and TCRdist3 normalized distances (between 0 and 1), each approach contributing to 50% of the final new TCR distances. The corresponding clustering tree (**SI FIG. 2**) presents 30 color changes and a pMHC-distance of 0.36, which is better than TCRdist3 but worse than TCRpcDist-3D, illustrating that there was no added value using the two approaches in this manner and in this particular case. We are working on combining our approach with TCRdist3 and other methods using several consensus approaches to see if TCR clustering efficiency can be improved.

We have quantified the outcome of the similarity principle using both approaches, and analysed the frequency at which the TCR with the smallest distance from a given one recognizes the same pMHC. We found that the two approaches are performing similarly. TCRdist3 more successfully paired TCRs with the same specificity when considering the TCR with closest distance (ranked 1), with a success rate of 46.7% compared to 44.4% for TCRpcDist-3D (**FIG. 2d**). However, TCRpcDist-3D better performed when 2 and 5 TCRs were considered (and not only one as above), with success rates of 64.4% and 75.6%, respectively, compared to 62.2% and 71.1% for TCRdist3. At rank 10, both approaches yield the same success rate. These results are in line with the lower scores for color changes and pMHC-distance exhibited by TCRpcDist-3D when analyzing the 45-TCRs dataset. Indeed, in this clustering, the metrics used compare each TCR with respect to a group of potentially the same specificity and not just with respect to a single TCR (rank1). Further comparisons between TCRpcDist-3D and TCRdist3 can be seen in **SI (PDB-set-84-SI.pdf)**.

### By applying distance thresholds on TCRpcDist-3D yields > 90% accurate pairing TCRs of the same specificity

Encouraged by these results, we have applied TCRpcDist-3D to a large and public data set of TCRs with known pMHC, the 10X dataset ( Single Cell Immune Profiling Dataset by Cell Ranger 3.0.2, 10x Genomics). We have worked with 8’528 TCRs (**SI Table 3**) for which we determined 3D structural models, calculated nSESA and computed the probability of a TCR pair to share the same pMHC as a function of the TCRpcDist-3D distance (**FIG. 3**). We have observed that at a distance of 0.15, the probability of a peptide pair to share the same pMHC is 50%. This probability increases when the distance between two TCRs decreases, reaching a maximum probability of 85% when the distance is 0. Of note, having 0 distance among a given TCR pair indicates an 85% likelihood that they share the same specificity.

Detailed analysis was performed without the KLGGALQAK viral peptide that represents 77.1% of the database (**SI Table 5** for the pMHC representation in the 10X set) and therefore could (1) bias the analysis performed, as the probability of randomly find a TCR recognizing the peptide KLGGALQAK is 60% and (2) make impossible to visualize details in the clustering tree. We re-computed for this subset the probability of a TCR pair to share the same pMHC as a function of the TCRpcDist-3D distance for the closest pair (**FIG. 3**) and the behavior is nearly the same. At a distance of 0.17 the probability of a peptide pair to share the same pMHC is 50%. Again, the lower the distance between two TCRs the higher the probability they share the same pMHC, reaching a maximum probability of ∼90% when the distance is 0. This public repertoire (SI Table 3 excluding KLGGALQAK) contains TCRs recognizing different pMHC (25 different peptides and 7 different alleles) characteristic of infectious diseases (19/25 pMHC) and cancer (6/25 pMHC), which can mimic experimental settings (**SI Table 2** and **SI table 6**).

We studied how frequently, starting from a given TCR, it is possible to find another TCR with the same specificity in the 1, 2, 5 and 10 top-ranked TCRs with closest distances (**FIG. 4**). We performed this analysis without any distance threshold and afterwards with thresholds of 0.15 and lower (**FIG. 4**) considering that the probability of sharing the same specificity is higher at these values (inversion in the sigmoid curve, **FIG. 3**). TCR pairs with the closest TCRpcDist (rank 1) share the same specificity in 53.9% of the cases, which is much better than what can be obtained by random assignments (19% success; p-value<0.0001). The predictive power of our TCRpcDist-3D is increased if we consider the top 5 or the top 10 closest TCRs to a given TCR. Indeed, the specificity of a given TCR is shared by at least one of the 5 and 10 top-ranked TCRs according to TCRpcDist-3D in 78.3% and 85.0% of the cases, respectively (p-value < 0.0001). The success can be further increased if a threshold of 0.15 for the closest pair is applied at the cost of decreasing the number of TCRs for which a comparison can be achieved. For example, in 91.0% of the cases, a given TCR of the dataset will share its specificity with at least one of the other 10 top-ranked TCRs according to TCRpcDist-3D, if we only consider cases where these TCRpcDist values were lower than 0.15 for the rank1. However, with this threshold, a comparison can be made for only 55.5% of the input TCRs. By applying more stringent thresholds we can increase significantly the predictive ability, at the cost of decreasing the number of TCRs that can be treated. For instance, in 97% of the cases, a given TCR will share its specificity with at least one of the 10 top-ranked other TCRs of the set, if we set a distance threshold lower than 0.025; Nevertheless, at this threshold, this analysis can only be made for 19.01% of the TCRs from the repertoire. We conclude that TCRpcDist lower than 0.15 seems to be a good compromise between accuracy and number of TCRs under study.

**FIG. 3.**
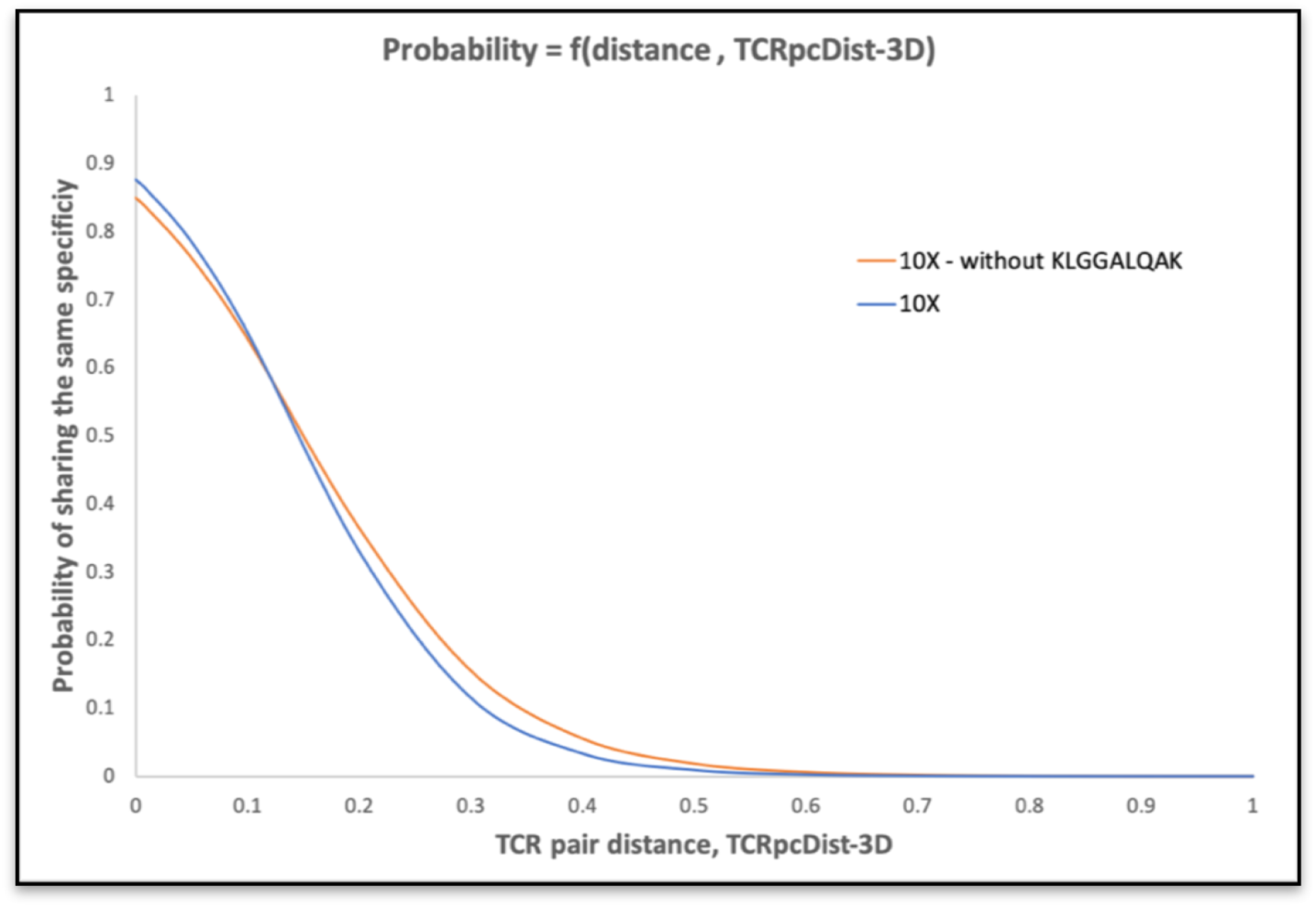
Probability of a TCR pair to share the same specificity as a function of their TCRpcDist-3D value. The sigmoid curve in blue was obtained using 8’528 TCRs with known specificities. The sigmoid curve in orange was obtained after removing the TCRs recognizing the overrepresented KLGGAQAK peptide.

**FIG. 4.**
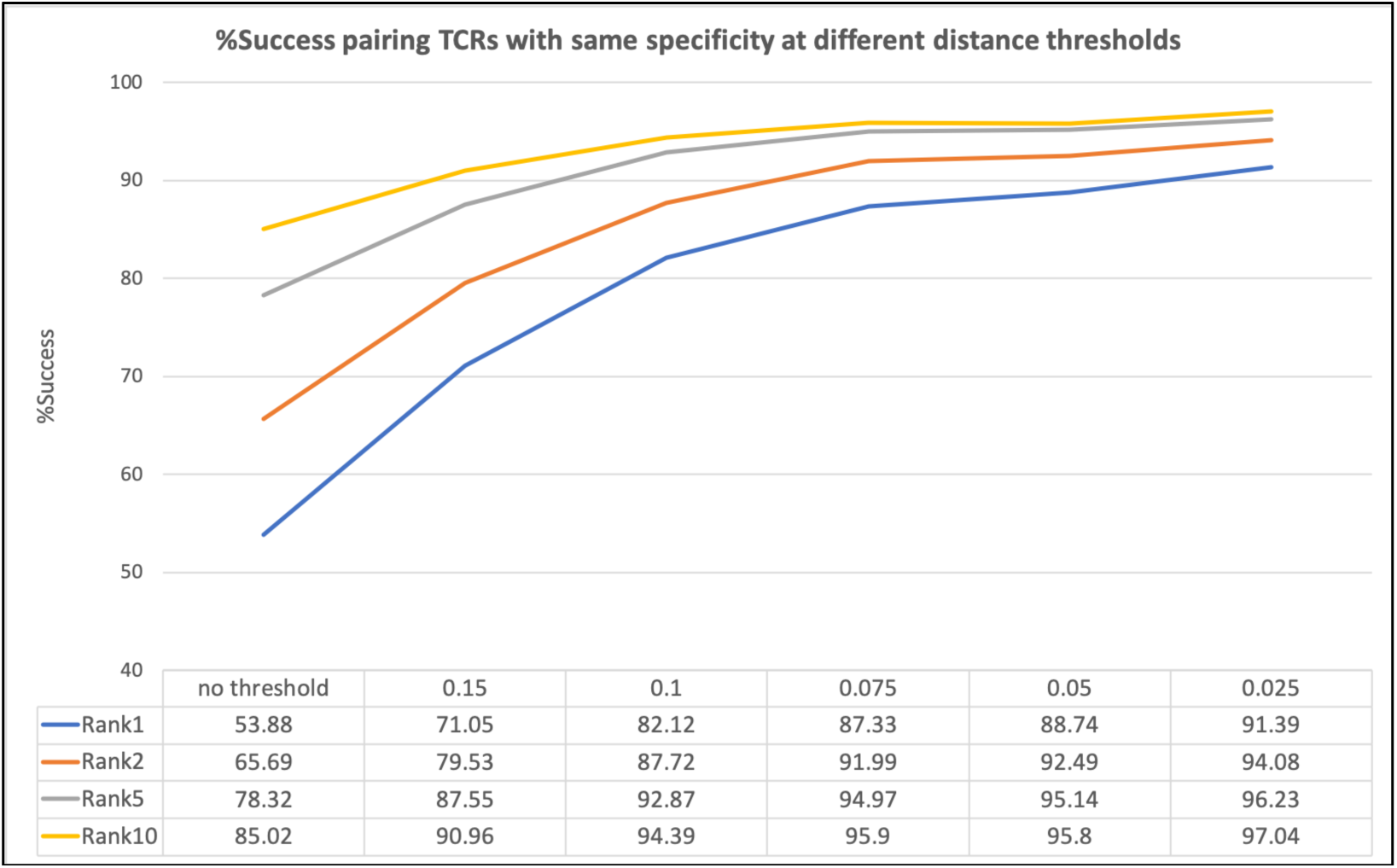
Percentage of success in pairing TCRs with the same specificity at different TCRpcDist-3D distance thresholds using the 10X Genomics dataset, after removing the overrepresented KLGGALQAK peptide.

We can notice that TCRs less frequently found in the reference set (<0.72%, **Table 2**) are rarely paired with the TCR of the same specificity, while TCRs with higher representations are easier to pair (**Table 2**). To tackle the problem of different abundance in the repertoire (which we decided to keep to mimic clinical data) we have introduced the enrichment factor (EF), a metric which determines if the abundance of the TCRs with a given pMHC specificity has increased in the top10 closest TCRs relative to the average abundance in the entire reference database. Indeed, the 10 top-ranked TCRs according to TCRpcDist-3D might accidentally contain TCRs with the above-mentioned specificities only because they are profusely represented in the repertoire. The analysis of the EF of the TCRs with some specificities allows to check if their presence in the top 10 is only an accidental result of their abundance in the repertoire, or an effect of their similarity to the input TCR, increasing the confidence in the prediction.

**Table 2.** Frequency of the TCRs with the same specificity correctly paired in the rank1 and in the rank10 when no threshold is applied. Results obtained when applying TCRpcDist-3D to the 10X Genomics data without the overrepresented peptide KLGGALQAK.

| Frequency | Peptide | Relative frequency<br>rank1 | Relative frequency<br>rank10 |
| --- | --- | --- | --- |
| 25.20 | AVFDRKSDAK | 34.08 | 93.10 |
| 0.72 | AYAQKIFKI | 0.00 | 0.00 |
| 0.10 | CLLWSFQTSA | 0.00 | 0.00 |
| 0.31 | CYTWNQMNL | 0.00 | 0.00 |
| 6.44 | ELAGIGILTV | 62.70 | 91.27 |
| 0.46 | FLASKIGRLV | 0.00 | 0.00 |
| 0.87 | FLYALALLL | 70.59 | 100.00 |
| 26.33 | GILGFVFTL | 84.66 | 96.12 |
| 2.25 | GLCTLVAML | 43.18 | 88.64 |
| 0.20 | IMDQVPFSV | 0.00 | 0.00 |
| 0.15 | IPSINVHHY | 0.00 | 0.00 |
| 6.85 | IVTDFSVIK | 32.84 | 64.93 |
| 0.46 | KTWGQYWQV | 0.00 | 0.00 |
| 0.15 | KVLEYVIKV | 0.00 | 0.00 |
| 0.31 | LLDFVRFMGV | 0.00 | 0.00 |
| 0.82 | LLFGYPVYV | 0.00 | 6.25 |
| 0.20 | MLDLQPETT | 0.00 | 0.00 |
| 0.15 | QPRAPIRPI | 0.00 | 0.00 |
| 21.17 | RAKFKQLL | 69.08 | 97.34 |
| 5.32 | RLRAEAQVK | 9.62 | 44.23 |
| 0.72 | RTLNAWVKV | 0.00 | 7.14 |
| 0.15 | SLFNTVATL | 0.00 | 0.00 |
| 0.36 | SLFNTVATLY | 0.00 | 0.00 |
| 0.15 | YLLEMLWRL | 0.00 | 0.00 |
| 0.15 | YLNDHLEPWI | 0.00 | 0.00 |

### Successful TCRs deorphanization of tumor-infiltrating lymphocytes from cancer patients using TCRpcDist-3D

TCRpcDist-3D and the clustering described herein were used for the prediction of TCR specificity, *i.e.*, predicting which pMHC a TCR could bind. This was done by applying the similarity principle which states that similar molecules are likely to share similar bioactivities. Following a strategy well-known in computer-aided drug design to predict the targets of bioactive compounds(*41, 42*), consisting in (i) reverse-screening a library of TCRs for which we know the specificity (a.k.a. the reference set) to identify those that are the most similar to the orphan TCR according to TCRpcDist-3D, and (ii) infer the orphan TCR’s probable specificity from the top 1, 2, 5 to 10 most similar TCRs in the reference set and their distance from the orphan TCR. Of course, using this approach, it is only possible to predict a potential specificity for an orphan TCR if it belongs to the specificities of the reference TCRs with which it will be compared. Consequently, due to the limited number of TCRs for which the cognate pMHC is already known, it may be that none of the orphan TCR binds any of the candidate epitopes.

We have studied 8’224 intratumoral orphan TCRs from four melanoma patients, 1856 from patient1 (0YM1), 2792 from patient2 (058C), 2011 from patient3 (13WU), and 1564 from patient4 (1AR4) (**SI Table 2**) and screened them against the TCRs with known specificity within the same patient for the first three patients and within all patients for the fourth (1AR4). The 45 TCRs of the reference set, along to the patients in which they were found and their specificities, are listed in **SI Table 6**. For patient 1AR4 we knew in advance that its orphans TCRs would react *vs* a pool of 32 pMHC (i.e. the CEF pool, with 32 well-defined viral epitopes from Cytomegalovirus, Epstein-Barr virus, and Influenza viruses, **SI Table 6**) and consequently we screened them against TCR specific to these peptides, even though they belong to a different patient. Restricting the screen of the orphan TCRs against TCRs with known specificity within the same patient for the first three patients limits the search space but guarantees that the corresponding pMHCs are effectively present in the patient in which the orphan TCR was found.

The analysis of all the possible pairs (CD8+ TCR orphan – CD8+ TCR with known specificity] showed that only 11 orphan TCRs have a TCRpcDist-3D lower than 0.15 to a given TCR with known specificity (**FIG. 5a**). We tested them experimentally together with 5 more orphan TCRs that present TCRpcDist-3D [CD8+ TCR orphan – CD8+ TCR with known specificity] distances between 0.15 and 0.25 to a reference TCR, with consequently lower probability of a correct specificity prediction (<<50%, **FIG. 3**). The predicted cognate pMHC of 2 TCRs out of these 16 orphan ones, were experimentally validated (**FIG. 6**). They correspond to those exhibiting the lowest distances together with the highest enrichment factor, i.e. a TCRpcDist-3D of 0.11 and 0.14 to the closest TCRs with known specificity and an enrichment factor of 4.5 in both cases(**FIG. 5a**). These two TCRs are specific to TPRVTGGGAM:HLA-B*07 and, in the set of 45 reference TCRs, spontaneously group with the TCR that recognizes TPRVTGGGAM:HLA-B*07 (**FIG. 5b**). The sequence details of the orphan TCRs experimentally validated and their closest TCR with known specificity can be seen in **Table 3**.

**FIG. 5.**
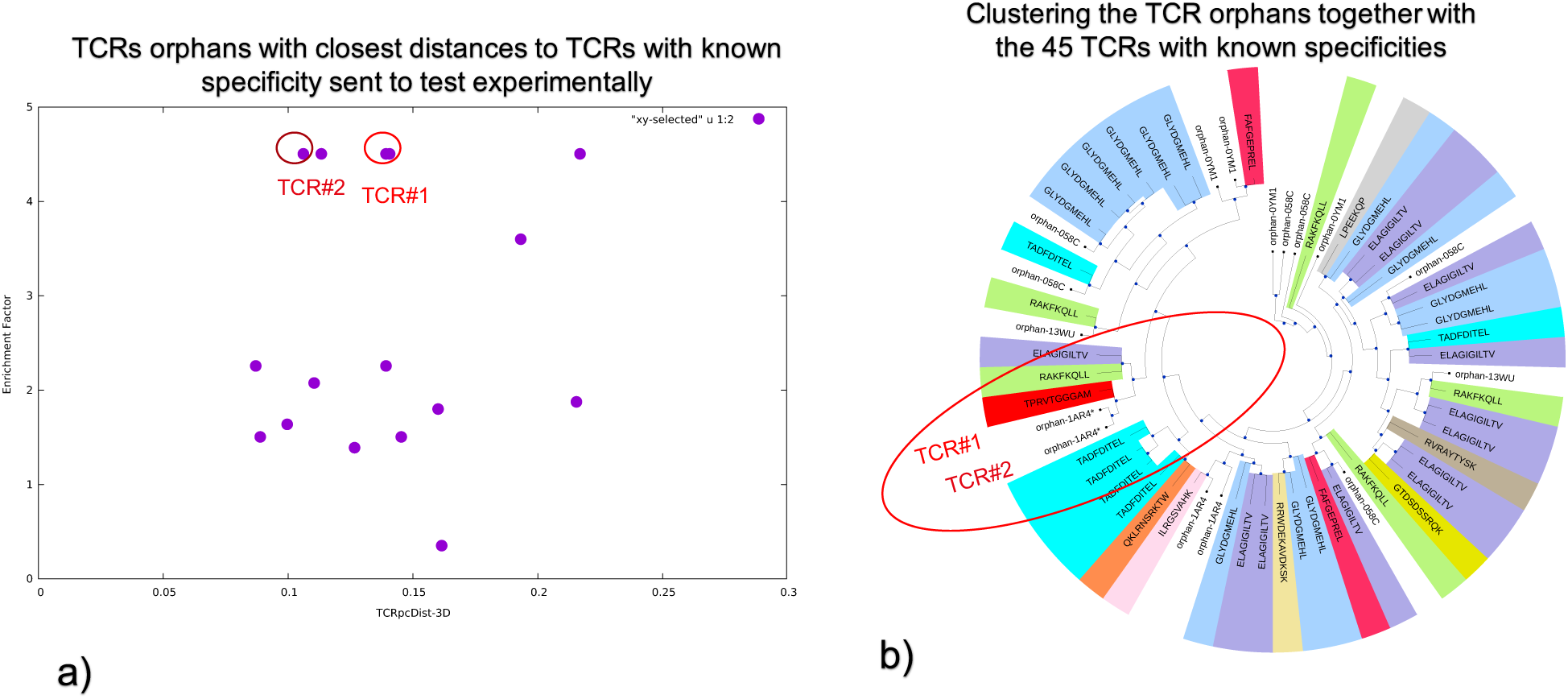
Using TCRpcDist-3D to deorphanize TCRs found in cancer cells of patients. a) TCRs orphans tested experimentally. Each dot corresponds to the TCRpcDist-3D value between the orphan TCR and the closest reference TCR, and the corresponding EF value. b) the clustering tree of the TCR orphans tested experimentally

**FIG. 6.**
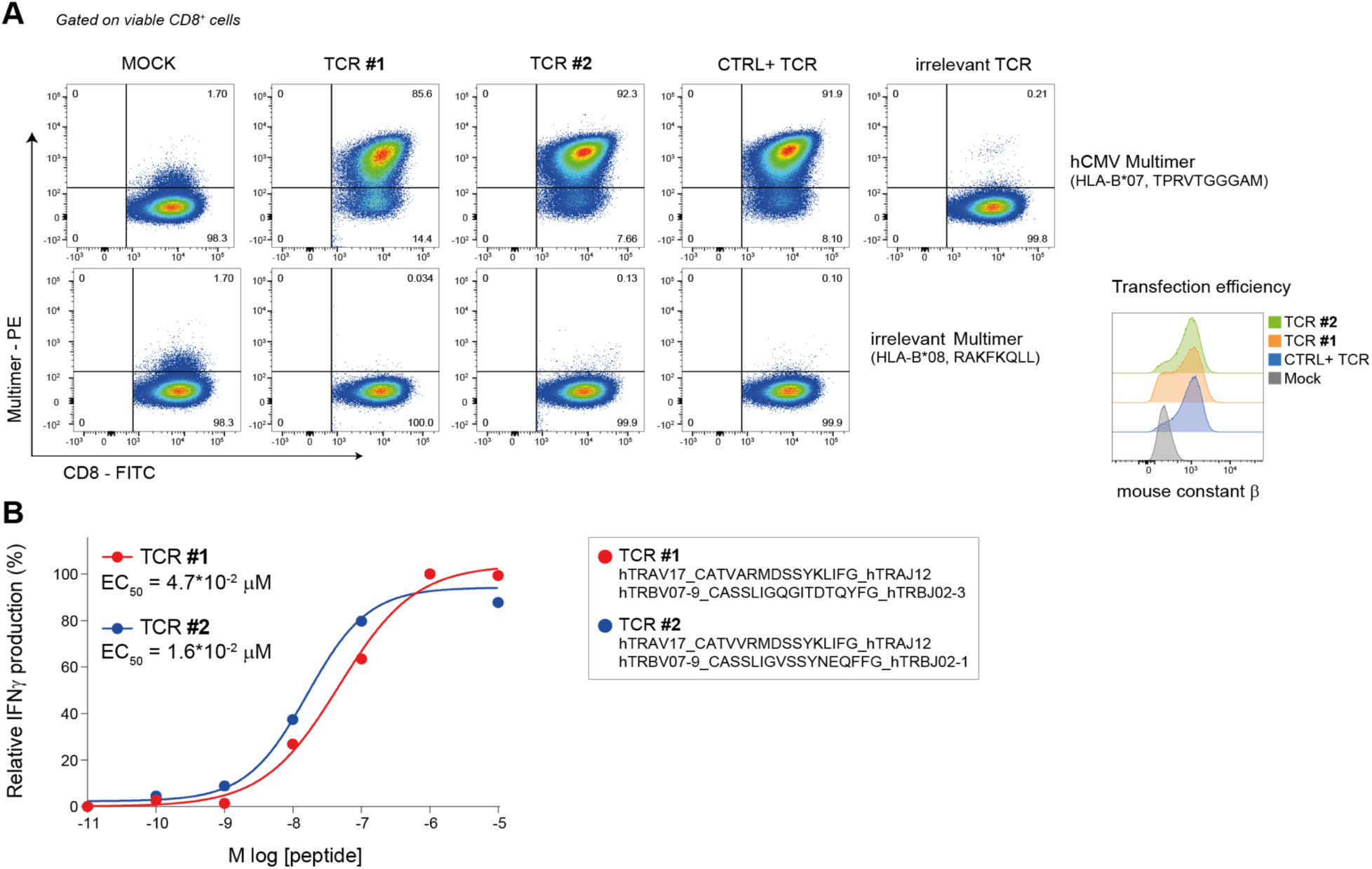
Validation of peptide-specificity and functional characterization of two TCRs predicted through TCRpcDist-3D. (A) Validation of antigen-specificity for two positive TCRs found by TCRpcDist-3D screening. TCRalpha- and TCRbeta-coding RNA was transfected into recipient Jurkat cells engineered for human CD8 expression and CRISPR TCRalphabeta-KO. After over-night incubation cells were stained with the CMV pp65-multimer (TPRVTGGGAM, HLA-B*07). A previously identified pp65-specific TCR and an irrelevant TCR were used respectively as technical positive (CTRL+) and negative control. An irrelevant EBV B2LF-1-multimer (RAKFKQLL, HLA-B*08) was used to further confirm the specificity of the two predicted TCRs. (B) The functional avidity of both CMV pp65-specific TCRs was measured using activated primary T cells. Shown are the normalized relative frequencies of IFNγ-producing T cells and the EC_50_ (effect concentration 50%, peptide concentration required for half-maximal T cell activation) is given for each TCR. Color-coded corresponding TCR sequences are reported on the right.

**Table 3.** Orphan TCRs for which the specificity was correctly predicted and their closest TCR with known specificity.

|  | TRAV | TRAJ | CDR3a | TRBV | TRBJ | CDR3b | peptide |
| --- | --- | --- | --- | --- | --- | --- | --- |
|  | TRAV17 | TRAJ12 | CATVVRMDSSYKLIFG | TRBV7-9 | TRBJ2-1 | CASSLVGEGWSDEQFFG | TPRVTGGGAM |
| TCR#2 | TRAV17 | TRAJ12 | CATVVRMDSSYKLIF | TRBV7-9 | TRBJ2-1 | CASSLIGVSSYNEQFF | orphan-1AR4 |
| TCR#1 | TRAV17 | TRAJ12 | CATVARMDSSYKLIF | TRBV7-9 | TRBJ2-3 | CASSLIGQGITDTQYF | orphan-1AR4 |

## DISCUSSION

We have presented a new definition of the distance between TCRs, TCRpcDist. The clustering pipeline consists of four main steps. First, all possible sliding windows of 4 residues that constitute the so-called 4-mer subunits are identified. The CDR residues that cannot contact directly the peptide, as determined by their solvent accessibility in the structural models, can be excluded from the process. Next, each 4-mer subunit is converted into a biophysicochemical representation using 5 Atchley factors. For each CDR of a pair of TCRs, all the *n* 4-mer motifs that are possible to construct from the first TCR with all the *m* possible 4-mer motifs of the second TCR are compared. This results in n×m matrix comparisons for each CDR for each pair of TCRs. The matrix comparisons are performed via a Manhattan distance score normalized over the maximum possible distance. This score ranges from 0, for 4-mers sharing exactly the same biophysicochemical properties, to 1, for 4-mers that have the highest difference in biophysicochemical properties. The method was developed using TCR:pMHC PDB structures before being tested for use with TCR homology models for broader applications. The clustering accuracy is found maximal when a weighting of about 30% is applied to the subset of amino acids in CDR3α or CDR3β and a weighting of 10% are given to the subset of amino acids in CDR1α, CDR2α, CDR1β, or CDR2β, respectively. The clustering accuracy is better when, together with the aforementioned weighting factors, only residues sufficiently exposed to the solvent, thus potentially able to contribute to the pMHC binding interface, are considered to be part of the 4-mers.

TCRpcDist-3D is a state-of-the-art approach that competes favorably with existing approaches as shown by comparison with TCRdist3. Similar to the method of Ostmeyer et al., TCRpcDist uses the Atchley factors rather than the sequences to calculate distances between TCRs. However, unlike that approach, but similarly to TCRdist, it takes CDR loops 1, 2 and 3 into account, and not just CDR3β. TCRpcDist differs however from TCRdist by the fact that the latter considers also CDR loop 2.5 and performs a global sequence alignment prior to scoring the latter using BLOSUM62 derived parameters, while TCRpcDist-3D tries to identify and uses the most relevant 4-mers (i.e. four consecutive residues) to account for the fact that not all residues are likely to bind directly to pMHC. To accentuate the focus on the residues most likely to interact with the pMHC, TCRpcDist-3D limits the search for 4-mers containing only sufficiently solvent exposed residues according to TCR structural models. TCRpcDist is therefore a molecular and biophysical approach that allows calculating distances between TCRs based on the physicochemical properties of the solvent exposed 4-mers that are possible to construct from CDR1α, CDR2α, CDR3α, CDR1β, CDR2β, and CDR3β sequences. Such distances can subsequently be used to cluster TCRs, or apply the similarity principle to try to predict TCR specificities. Moreover, TCRpcDist is broadly applicable and can be generalized without the need for supervised training.

We have challenged our approach and predicted specificities for two orphans from a pool of 8’224 TCRs. Among the 11 predictions for which the similarity principle could be applied with some confidence (distance < 0.15) we have validated the specificity for two orphans for the same viral peptide, the ones with higher EF. Unfortunately, we could not predict and validate a TCR orphan binding a neo-antigen. This might become achievable in the future, when more TCRs binding known neoantigens will be available. Applying TCRpcDist-3D to additional private data with known and unknow specificities, from more patients and other cancer types, is ongoing.

TCRpcDist-3D and the clustering described herein can be used to analyze the diversity of TCR repertoires, by grouping together those likely to bind the same pMHC. In addition, it can also be used for the prediction of TCR specificity, *i.e.*, predicting which pMHC a TCR could bind. One possible approach is to cluster TCRs for which the cognate pMHC is known, together with orphans TCRs for which the target is unknown. The positioning of these orphan TCRs indeed provides an indication about possible cognate pMHC, *i.e.* those belonging to a group of TCRs that recognize one given peptide or those that have the closest distance to a TCR with a given specificity. The capacity of TCRpcDist-3D to predict specificities for orphan TCRs in cancer patients was validated experimentally. TCRpcDist is able to provide new insights into T cell responses captured in the TCR repertoire and will facilitate the development of new clinical strategies to treat and monitor not only cancer but other infectious diseases.

### Limitations of the study

Our approach demonstrates high clustering efficiency when compared with the competitors but there is still room for further improvements. The fact that the TCRpcDist calculates TCR distances based on 4-mer features is a limitation, particularly when working with TCRs with long CDR loops, where more than 4 residues can be involved in the binding of the antigen. Using sequence features longer than 4-mer in the approach may improve the clustering efficiency for longer CDR loops but will not be applicable for short ones. We are exploring the feasibility of including features of variable lengths. Another limitation of TCRpcDist is the fact that it uses one single conformation per TCR model, i.e. the lowest energy one among 10 conformations, to calculate the residues that may interact with the pMHC. The dynamic behavior of the CDR loops and the stochastic nature of the modelling approach may result in different residues available to recognize the pMHC within the same TCR sequence if we model it in different times. To circumvent this issue, we could sample more conformations to find the low-energy one and/or make use of a conformational ensemble in order to determine the residues able to interact with a pMHC. Of note, although more sampling and an approach that makes use of a conformational ensemble would be feasible, it would also be substantially more time consuming which could be a limitation for large scale applications, for example on thousands of orphan TCRs from clinical sets.

Regarding the specificity predictions, TCRpcDist proved to be able to fish and deorphanize TCRs in 4 cancer patients under clinical investigation. Additional specificity predictions in these and other patients will further be essential to describe the impact of our approach in cancer immunotherapy. Further studies are ongoing thanks to a larger private data set with more patients and more TCRs with known specificities.

To further improve our clustering efficiency and deorphanization capability we are also working on (i) combining TCRpcDist with other methods following a consensus approach, so that we can combine their individual strengths and on (ii) applying to additional databases.

## MATERIALS AND METHODS

### TCRs datasets

Our development set consists in of 54 CD8+ TCR structures with known pMHC (**SI Table1**) taken from a set of 151 TCR:pMHC complexes whose experimental structure were retrieved from the Protein Data Bank in June 2020 (*33*). TCR duplicates (having exactly the same CDR 1, 2, 3; α and β composition) and singleton pMHC (pMHC recognized by only one single TCR, thus preventing the possibility of cognate TCR comparison) were removed from this set. Human and mouse structures were considered to increase the sampling size.

Our assessment set consists in a private collection of 45 CD8+ TCR sequences with known pMHC (**SI Table2**) found in 4 melanoma cancer patients. Further details about the patient data and TCR determination for these cancer patients are given below across this methods section.

Further assessment of our approach was done on a much larger data set that consists in CD8+ TCR sequences taken 10X Genomics data (10X Genomics, version 3.0.2). Only true cells (with a “True” label in the “is_cell” column of the all_contig_annotations.csv file) and cells expressing high confidence were kept for further analyses. TCR pairs lacking some important information like one of the chains, one gene, the CDR3 information and the cognate pMHC were also removed. Moreover, we have worked with TCRs that express one single α and one single β chains and removed redundant, singleton and cross reactive TCRs. This generated a list of 8’528 CD8+ TCRs (**SI Table3**) that could be modelled in 3D (see below) and of which 77.1% bind the CMV peptide KLGGALQAK bound to the A*03:01.

Finally, 8’224 orphan TCRs (**SI Table 2)** determined by single cell experiments and found in 4 cancer patients were used with the objective of deorphanizing some of them. Further details about the patient data and TCR determination for these cancer patients are given below.

### Modelling TCR 3D-structures from sequence

The Rosetta “TCRmodel” protocol (*28*) was applied to find the best TCR templates and model the 3D structures of the TCRs from the sequences (10X Genomics and private sets). A total of 10 models were produced for each TCR, and the highest ranked one according to the Rosetta energy function (*28*) was selected as the final model for the TCR 3D-structure. The number of TCRs sequences previously mentioned in the section *TCR datasets* correspond to TCRs for which 3D models could be obtained. Indeed, 19.6% of TCR sequences from 10X Genomics could not be modelled due to the lack of relevant templates.

### Solvent Accessibility calculation

The solvent accessibility of each CDR1α, CDR2α, CDR3α and CDR1β, CDR2β and CDR3β residue was determined as the relative solvent excluded surface area (SESA) computed with the MSMS package of the UCSF Chimera software (*43, 44*), as described elsewhere (*45*). The normalized SESA, nSESA, was calculated by normalizing the surface area of the residue in the TCR of interest by its surface area in a reference state. The latter was defined as the Gly-X-Gly tripeptides in which X is the residue type of interest (*46*). nSESA thus ranges from 0% for totally buried residues to 100% for residues with the side chain fully exposed to the solvent.

### TCRs distance calculation and clustering

The clustering pipeline is summarized in **FIG. 7** and consists in the following main steps:

**FIG. 7.**
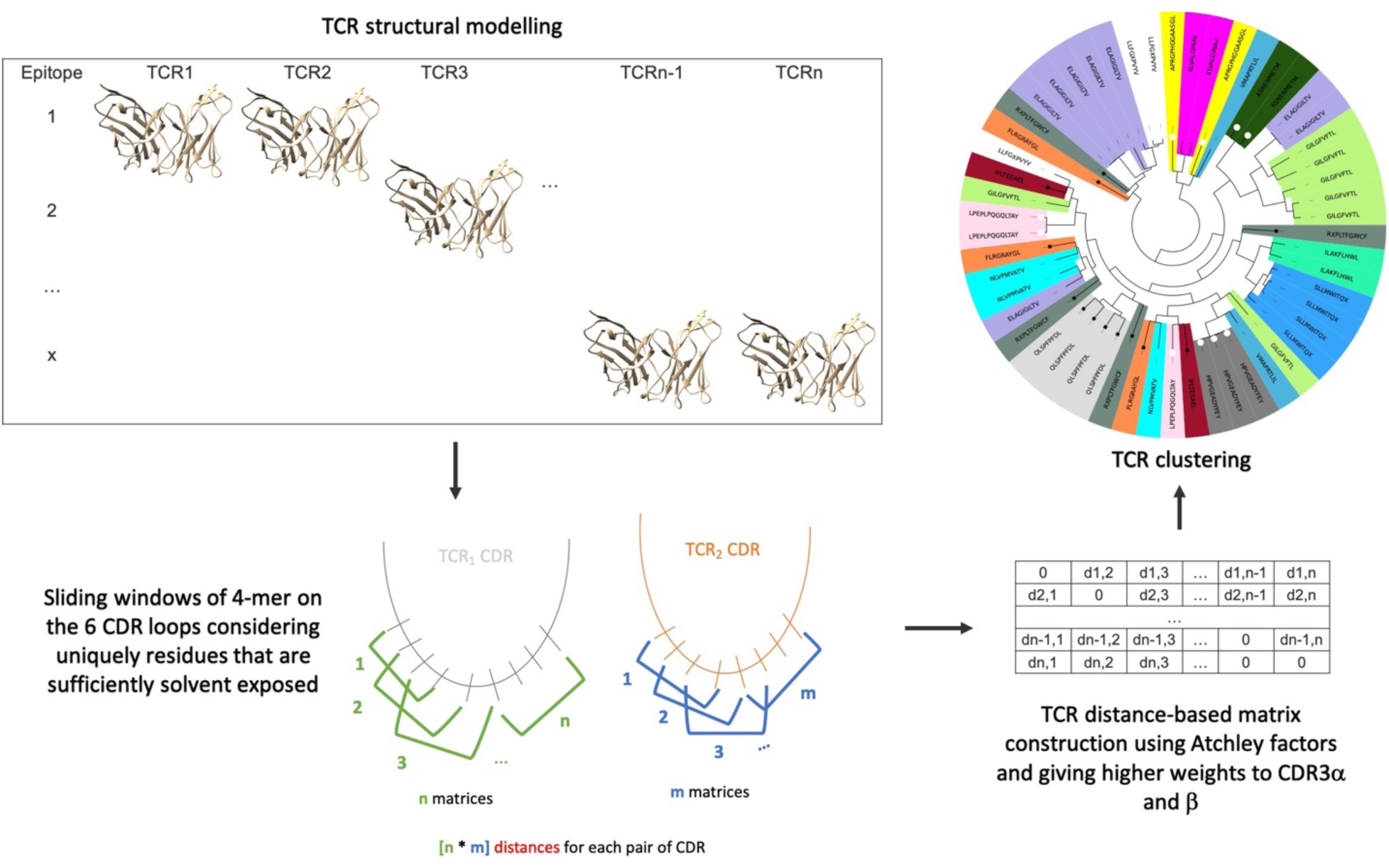
Representative scheme for the clustering pipeline used in TCRpcDist-3D.

First, all possible sliding windows of 4 residues that constitute the so-called 4-mer subunits are identified. The first four and last 3 residues from the CDR3 are excluded from the process. Alternatively, CDR residues with low SESA in the structural models may be excluded from this process since these residues are unlikely to contact the pMHC. Indeed, after a benchmark of the TCR clustering of the PDB set (details in the results section), using only the residues of CDR1s and CDR2s (α and β) with SESA > 5% and CDR3s (α and β) with SESA>20% was found to improve the quality of the clustering approach. Solvent accessibility per residue per TCR for the PDB set can be read in Supporting Information ( **SESA-PDB-54-allCDRs.tar.gz**).

Second, each 4-mer subunit was converted into a biophysicochemical representation using 5 Atchley factors that describe i) hydrophobicity, ii) secondary structure, iii) size/mass, iv) codon degeneracy, and v) electric charge (*26*). Given the encoding of residues using Atchley factors, we calculate the distance between two sets of 4 consecutive residues as the Manhattan distance between the two corresponding matrices M1 and M2, *i.e.*, d(M1, M2) where:

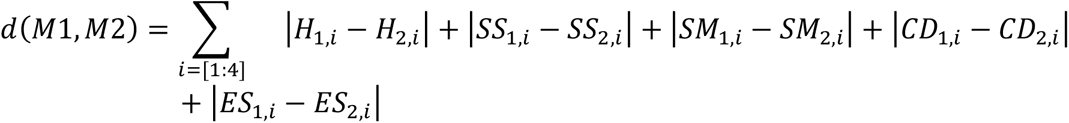

Here, H_1,i_ is the Hydrophobicity Atchley factor of residue *i* in Matrix 1, H_2,i_ is the Hydrophobicity Atchley factor of residue *i* in Matrix 1, SS_1,i_ is the Secondary Structure Propensity Atchley factor of residue *i* in Matrix 1, etc.

Third, to calculate the distance between two corresponding loops, such as the CDR3β of two different TCRs, the matrices of Atchley factors for the *n* possible 4-residues sliding windows of the first TCR and the *m* 4-residues sliding windows of the second TCR were generated. Then, the distance between each possible corresponding pair of matrices is comprehensively calculated. The smallest distance, d_min_, between all possible pairs of matrices is retained as the distance between the two CDR3β loops.

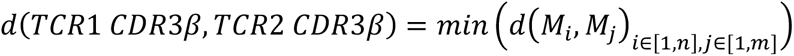

The above equation to calculate the distance between two CDR3β of two different TCRs is also applied to all CDR1s and CDR2s. Finally, the distance between the two TCRs is defined as the (possibly weighted) sum of the calculated distances between each pair of CDR1α, CDR2α, CDR3α, CDR1β, CDR2β, and CDR3β loops.

Of note, when analysing a set of multiple TCRs, all distances between each possible pair of TCRs are calculated as described above. These distances are then normalized by dividing each of them by the largest calculated distance for all possible pairs. Consequently, the final distance between two TCRs ranges from 0 (for two TCRs bearing an identical set of 4 residues on each considered loop – only CDR3β or all CDRs) to 1 (for two totally unrelated TCRs).

Finally, the calculated distances between TCRs within a set can be used to construct a distance tree. (**FIG. 7**). After an optimization of the TCR clustering approach using the PDB set (details in the results section), we found that the best clustering was obtained when giving a weight of 10% to the contributions of CDR1s and CDR2s (α and β) and of 30% to those of CDR3s (α and β) in the calculation of the distance. The generic hierarchical clustering algorithm UPGMA (unweighted pair group method with arithmetic mean) was used. The ETE3 python toolkit (*47*) was employed for the visualization of the clustering trees. Each TCR in the clustering tree is colored according to its specificity (pMHC). As a measure of the quality of the clustering, we determined the number of times the color changed between two successive nodes, starting from the upper node of the hierarchical clustering and turning clockwise. The quality of the clustering was also determined by a more quantitative metric, pMHC-distance, using the python library for phylogenetic computing, DendroPy (*48*), version 4.5.2 and the class PhylogeneticDistanceMatrix. pMHC-distance was defined as the average branch length distance between all possible pairs of TCR nodes that recognize the same pMHC.

### Patients and regulatory issues

Patients with stage III/IV metastatic melanoma, who had received several lines of chemotherapy and immunotherapy were included. Patients were enrolled in a single-center phase I trial of TIL-ACT (ClinicalTrials.gov NCT03475134, https://www.biorxiv.org/content/10.1101/2022.12.23.519261v1) under protocols approved by the respective institutional regulatory committees at Lausanne university hospital (CHUV), Switzerland. Patients’ recruitment, and study procedures were approved by regulatory authorities and all patients signed written informed consents.

### Tissue processing

Resected baseline tumors (prior TIL-ACT, tumors used to generate TIL products) were chopped into 1-2 mm2 pieces and cryopreserved in 90% human serum + 10% dimethyl sulfoxide (DMSO). For single-cell experiments, both frozen and fresh material were used as starting material. The day of the assay, pathofrozen pieces were thawed in RPMI + 10% FBS and chopped in small pieces using a scalpel. Tissue was dissociated in RPMI + 2% Gelatin (#G7041, Sigma-Aldrich) + 200 IU/mL Collagenase I (#17100-017, ThermoFisher Scientific) + 400 IU/mL Collagenase IV (#17104-019, ThermoFisher Scientific) + 5 IU/mL Deoxyribonuclease I (#D4527, Sigma-Aldrich) + 0.1% RNasin Plus RNase Inhibitor (#N2618, Promega) for 15-30min (depending of sample size and consistency) at 37°C and shaken at 160rpm. Digested cells were filtered using a 70 µm strainer and resuspended in PBS + 1% Gelatin + 0.1% RNasin. Cells were manually counted with hematocymeter then stained for viability with 50uM/mL of Calcein AM (#C3099, Thermo Fisher Scientific) and FcR blocked (#130-059-901, Miltenyi Biotec) for 15min at RT. After incubation and washing, cells were stained with CD45-APC (#304012, BioLegend) for 20min at 4°C. After washing, cells were resuspended in PBS + 0.04% BSA (Sigma-Aldrich) + 0.1% RNasin and DAPI staining (Invitrogen) was performed

### Single-cell RNA and TCR sequencing

CD45 live cells were sorted with the MoFlo AstriosEQ (Beckman Coulter) and manually counted to assess viability with Trypan blue. *Ex vivo* CD45 cells from tumors were resuspended at a density of 600-1200 cells µL^-1^ with a viability of >90% and subjected to a 10X Chromium instrument for single-cell analysis. The standard protocol of 10X Genomics was followed and the reagents for the Chromium Single Cell 5’ Library and V(D)J library (v1.0 Chemistry) were used. 12’200 cells were loaded per sample, with the targeted cell recovery of 7’000 cells according to the protocol. Using a microfluidic technology, single-cell were captured and lysed, mRNA was reverse transcribed to barcoded cDNA using the provided reagents (10X Genomics). 14 PCR cycles were used to amplify cDNA and the final material was divided into two fractions: first fraction was target-enriched for TCRs and V(D)J library was obtained according to manufacturer protocol (10X Genomics). Barcoded V(D)J libraries were pooled and sequenced by an Illumina HiSeq 2500 Sequencer. The second fraction was processed for 5’ gene expression library following the manufacturer’s instruction (10X Genomics). Barcoded samples were pooled and sequenced by an Illumina HiSeq 4000 sequencer.

The scRNA-seq reads were aligned to the GRCh38 reference genome and quantified using *Cellranger* count (10x Genomics, version 3.0.2). Filtered gene-barcode matrices that contained only barcodes with unique molecular identifier (UMI) counts that passed the threshold for cell detection were used for further analysis. The number of genes per cell averaged 1’240 (median: 1’458) and the number of unique transcripts per cell averaged 4’206 (median: 2’926). We obtained a total of 67’133 cells. Low quality cells exhibiting more than 10% of mitochondrial reads were discarded from the analysis, resulting in a final set of 61’1991 cells. The data was processed using the *Seurat R* package (version 3.2.2) as follows briefly: counts were log-normalized using the *NormalizeData* function and then scaled using the *ScaleData* function by regressing the mitochondrial, ribosomal contents and S phase and G2/M phase scores. Dimensionality reduction was performed using the standard *Seurat* workflow by principal component analysis followed by tSNE and UMAP projection (using the first 75 PCs). The k-nearest neighbors of each cell were found using the *FindNeighbors* function run on the first 75 PCs, and followed by clustering at several resolutions using the *FindClusters* function. Cells were annotated by looking at expression of the canonical *PTPRC* and *CD3E* markers where all clusters where found to be T-cells. The cells were then classified as CD8-positive, CD4-positive, double-negative (DN), double-positive (DP) and Tγδ as follows: cells with non-null expression of *CD8A* and null expression of *CD4* were defined as CD8-positive (and *vice-versa* for CD4-positive). Cells showing non-null expression of both genes were classified as DP. Due to notorious dropout events in single-cell data, cells lacking the expression of both markers were classified as follows: if a cell belongs to a cluster (taking a fine resolution of 10) in which the 75^th^ percentile expression of CD8 was higher than its 75^th^ percentile expression of CD4, it was classified as CD8-positive (and *vice-versa* for CD4-positive cells). If the 75^th^ percentile expressions of both markers equal 0, the cells were classified as DN. Finally, cells with an average expression scores of all *TRG* and *TRD*-related genes higher than 0.3 were assigned to be Tγδ cells. (https://www.biorxiv.org/content/10.1101/2022.12.23.519261v1) scTCR-seq (VDJ) data were aligned to the same human genome using the cellranger vdj (10X Genomics, version 3.1.0). Only true cells (with a “True” label in the “is_cell” column of the all_contig_annotations.csv file) were kept for further analyses. Cells from the VDJ sequencing were mapped to the scRNA-Seq data (5′GEX). This allowed to only select CD8^+^ TCR clones for downstream analyses.

### Peptide synthesis

Peptides produced by the Peptides and Tetramers Core Facility (PTCF) of the University of Lausanne were HPLC purified (≥90% pure), verified by mass spectrometry and kept lyophilized at −80°C.

### TCR validation

To validate antigen specificity and interrogate T cell reactivity, TCRα/β pairs were cloned into recipient activated peripheral T cells and Jurkat cells (TCR/CD3 stably transduced with human CD8α/β and TCRα/β CRISPR-KO), as previously described (*40*). In brief, paired α and β chains were annotated based on TCRpcDist and corresponding full-length codon-optimized DNA sequences were synthesized at GeneArt (Thermo Fisher Scientific) or Telesis Bio DNA served as template for in vitro transcription (IVT) and polyadenylation of RNA molecules as per the manufacturer’s instructions (Thermo Fisher Scientific), followed co-transfection into recipient T cells.

For the avidity assay, autologous PBMCs were resuspended at 10^6^ cells mL^-1^ in R8 medium supplemented with 50 IU mL^-1^ IL-2 (Proleukin). T cells were activated with Dynabeads Human T Activator CD3/CD28 beads (Thermo Fisher Scientific) at a ratio of 0.75 beads: 1 total PBMCs. After 3 days, beads were removed and activated T cells cultured for two extra days before electroporation or freezing. For RNA transfection, the Neon electroporation system (Thermo Fisher Scientific) was used. Cells were resuspended at 15-20×10^6^ cells mL^-1^ in buffer R and mixed with 500 µg of TCRα chain RNA together with 500 µg of TCRβ chain RNA and electroporated with the following parameters: 1600V, 10ms, 3 pulses.

The functional avidity of hCMV pp65-specific T cells was assessed by IFN-γ Enzyme-Linked ImmunoSpot (ELISpot, Mabtech) assay with limiting peptide dilutions (ranging from 10 µg mL^-1^ to 0.1 pg mL^-1^) as described(*49*). Briefly, 2×10^3^ transfected T cells were plated per well in a pre-coated 96-well ELISpot plate (Mabtech) together with 5×10^4^ PHA-activated autologous CD4 T cells as antigen-presenting cells and challenged with the specific peptide. EC_50_ values were derived by dose-response curve analysis (log(peptide concentration) versus response) using GraphPad Prism software (v.7, GraphPad). The peptide concentration required to achieve a half-maximal cytokine response (EC_50_) was determined and referred to as the functional avidity.

Jurkat cells were electroporated using the Neon electroporation system (Thermo Fisher Scientific) as described above for activated PBMCs and with the following parameters: 1,325 V, 10 ms, three pulses. After over-night incubation, electroporated Jurkat cells were interrogated by pMHC-multimer staining with the following surface panel: anti-CD3 APC Fire 50 (SK7, Biolegend), anti-CD8 Pacific Blue™ (RPA-T8, BD Biosciences), anti-mouse TCRβ-constant APC (H57-597, Thermo Fisher Scientific) and with viability dye Aqua (Thermo Fisher Scientific).

## Supporting information

SI-SESA-PDB-54-allCDRs.tar.gz

PDB-set-84-SI.pdf

SI-VariabilityOfOutcome

## Acknowledgments

We thank Drs Denarda Langaj Laniti and David Barras for their valuable help on single-cell data analysis of the four melanoma patients, as well as Rémy Pétremand, Aymeric Auger and Baptiste Murgues for their technical support.

## Author contributions

Conceptualization, MASP and VZ;

Methodology, MASP and VZ (computational) and JC, SB, MA and AH (experimental);

Software, MASP and VZ;

Investigation, MASP, JC, SB, FMR, GC, AH and VZ;

Data Curation, MASP and FMR

Writing – Original Draft, MASP and VZ

Writing – Review & Editing, MASP, JC, SB, FMR, GC, AH and VZ;

Visualization, MASP and VZ;

Supervision, AH and VZ;

Funding Acquisition, VZ, GC and AH

## Supplementary Materials

**SI Figure 1**. The 3D structure of the complex TCR:pMHC PDB ID 4JRX. Representation of TCR alpha chain in brown ribbon, TCR beta chain in blue ribbon with the CDR3β residues in stick and colored by atom with carbon atoms in blue, peptide in pink ribbon and MHC in purple ribbon. CDR3β amino acids are labeled in black (single letter code). It is show that the Pro (P), the 4rd residue in the CDR3β sequence, it is in contact with the bulged portion of the peptide.

**SI Table 1**. Training set of 54 TCRs with known pMHC taken from Protein Data Bank as of June 2020. Redundant TCRs and singleton TCRs were removed from an initial set of 151 TCRs.

Solvent Accessibility per residue per TCR for the set of 54 TCRs (SI Table 1) can be read in **SESA-PDB-54-allCDRs.tar.gz.**

**SI Table 2.** Private set of 45 TCRs with known specificity and 8’224 orphan TCRs from 4 melanoma patients. Some TCRs already disclosed in Arnaud *et al*. (*40*)and patients described in Barras *et al* (https://www.biorxiv.org/content/10.1101/2022.12.23.519261v1).

**SI Table 3**. Set of 8528 CD8+ TCRs with known pMHC taken from 10X genomics version 3.0.2.

**SI Table 4**. Sequence variability expressed by TCRs recognizing ELAGIGILTV in the PDB set and in the private set. For the full set of TCRs within each set report to SI tables 2 and 1, respectively.

**SI Table 5**. TCRs frequency for each one of the pMHC in the set of 8528 TCRs taken from 10X genomics (SI Table 3)

**SI Figure 2**. Hierarchical clustering of a set of 45 private TCRs (SI Table 2) recognizing 12 different pMHC combining TCRpcDist-3D and TCRdist3 normalized distances (between 0 and 1), each approach contributing to 50% of the final new TCR distances.

**SI Table 6**. TCRpMHC per patient among the 4 melanoma patients studied (one sheet per patient in the excel file). Per patient is given the allele and if the TCRs are tumor reactive or not.

**PDB-set-84-Si.pdf** – comparison between TCRpcDist-3D and TCRdist3 using a non-redundant PDB set of 84 TCRs.

**SI-SESA-PDB-54-allCDRs** – solvent accessibility calculations for the CDR loops of the PDB structures.

**SI-VariabilityofOuctome.xlsx** – Three independent TCRpcDist-3D distance calculations for the private set of 45 TCRs (sheet1, sheet2 and sheet3) and their standard deviations SD (sheet4). We observe that among the three independent runs there are only 94 TCR pairs distances with SD>0.1 and the maximal SD is 0.19. This indicates that only 47 pairs of TCRs among the 990 possible pairs report SD>0.1, i.e. 4.7%. The average SD over all the pairs is only 0.04, indicating that the variability on calculated TCRpcDist-3D distances between several runs, starting from the same input, remains very limited and does not change the main conclusions of TCR distance analysis.

