## Supplementary figures and images for "TCRpcDist: Estimating TCR physico-chemical similarity to analyze repertoires and predict specificities"

### Supplemental Figure 1

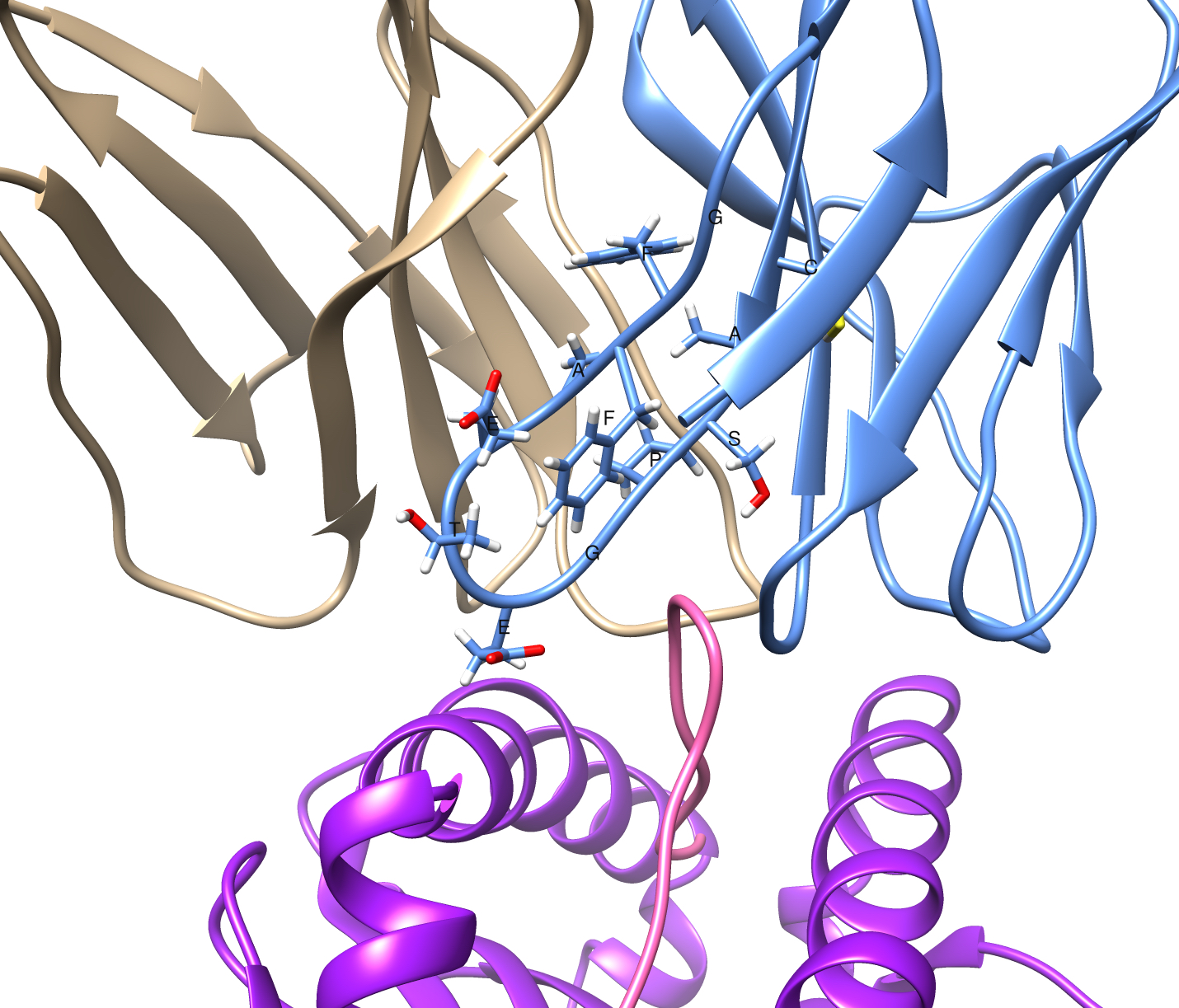

### Supplemental Figure 2

#### Clustering pMHC

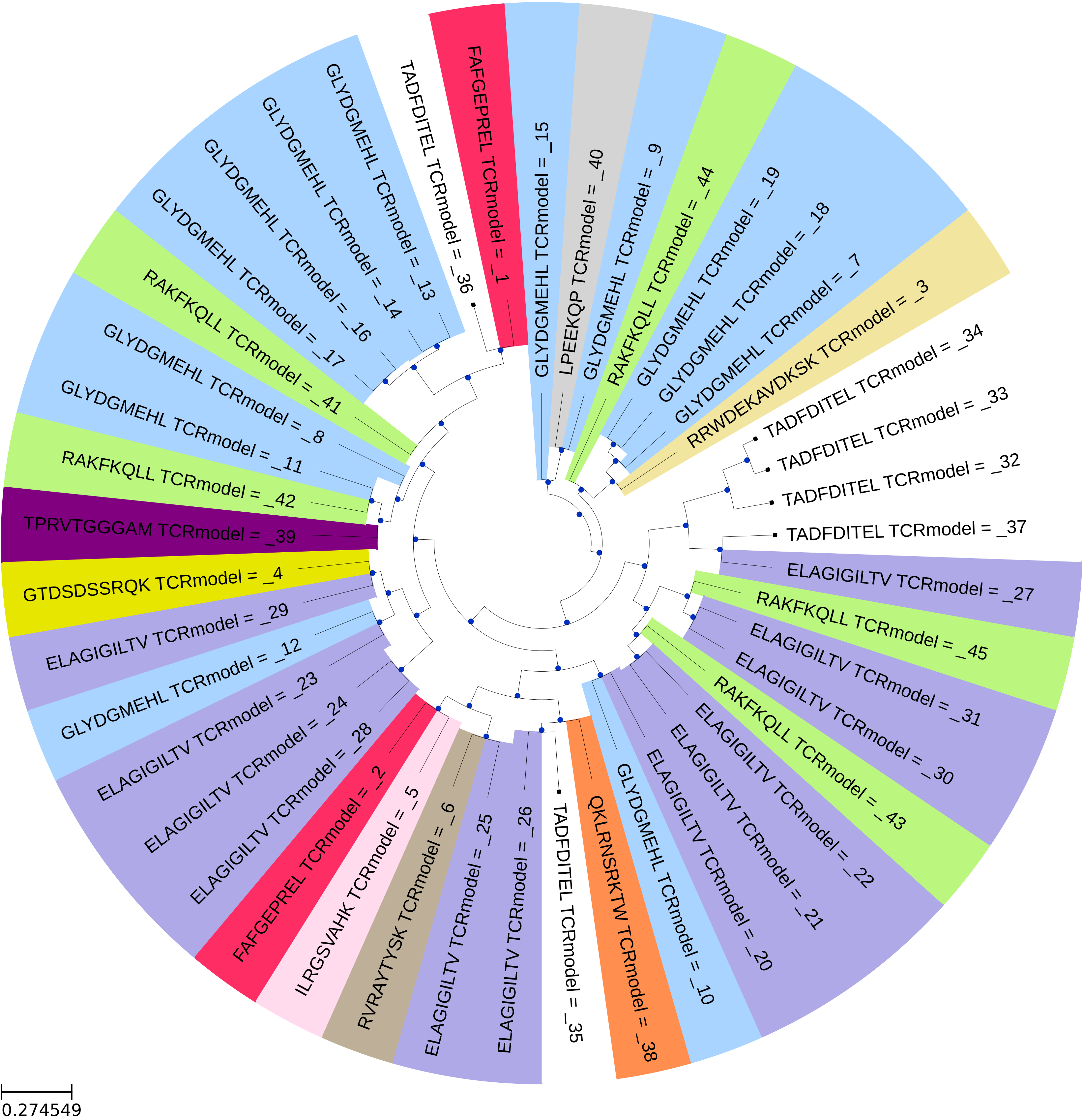
