## Supplementary material for "TCRpcDist: Estimating TCR physico-chemical similarity to analyze repertoires and predict specificities": PDB-set-84-SI.pdf

For purposes of comparison with competing approaches we have worked with a subset of 84 non-redundant and non-singleton CD8+ TCRs with known specificity taken from PDB (January 2023). We could not compare the initial developmental set of 54 TCRs from PDB with the TCRdist3 approach as it contains mouse and human TCRs together and we cannot input both species in the tcrdist3 algorithm. This time TCRpcDist-3D show better clustering efficiency and equal or higher accuracy for all the ranks.

### Set of 84 – human – no redundant (no single point mutation) – no singleton

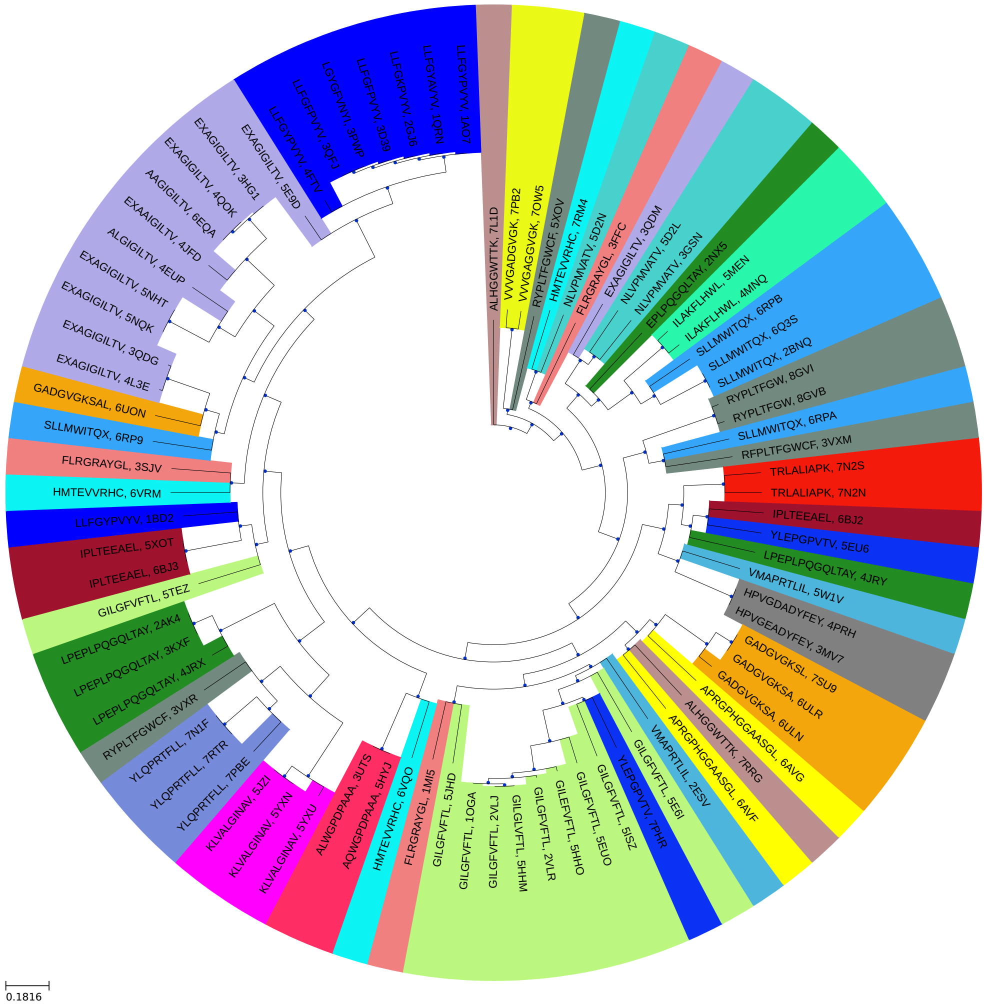


**TCRdist**
Color change = 44
pMHC-distance =0.40

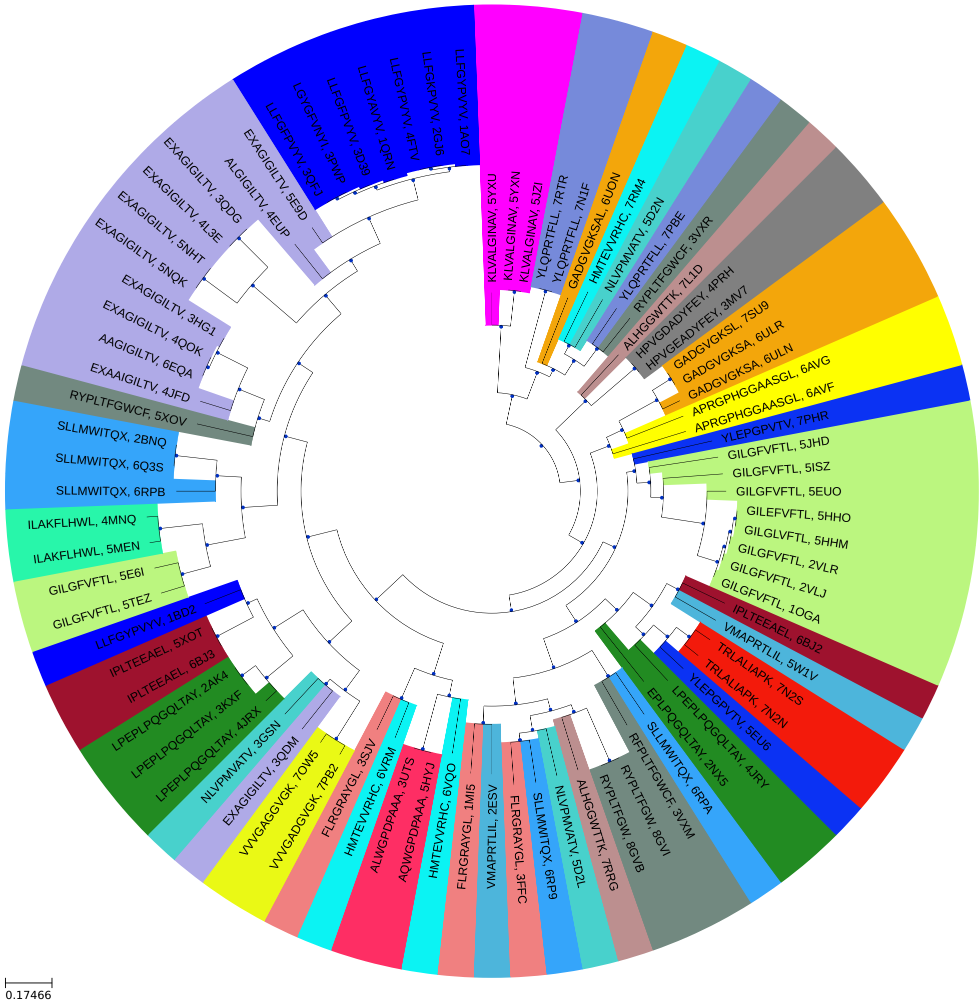


**TCRpcDist-3D**
Color change = 42
pMHC-distance =0.30
TCRs recognizing the same peptide are closer in the Tree

✓ TCRpcDist-3D has lower color change and lower pMHC-distance

### Distribution of the normalized distances in TCRpcDist-3D and in TCRdist3

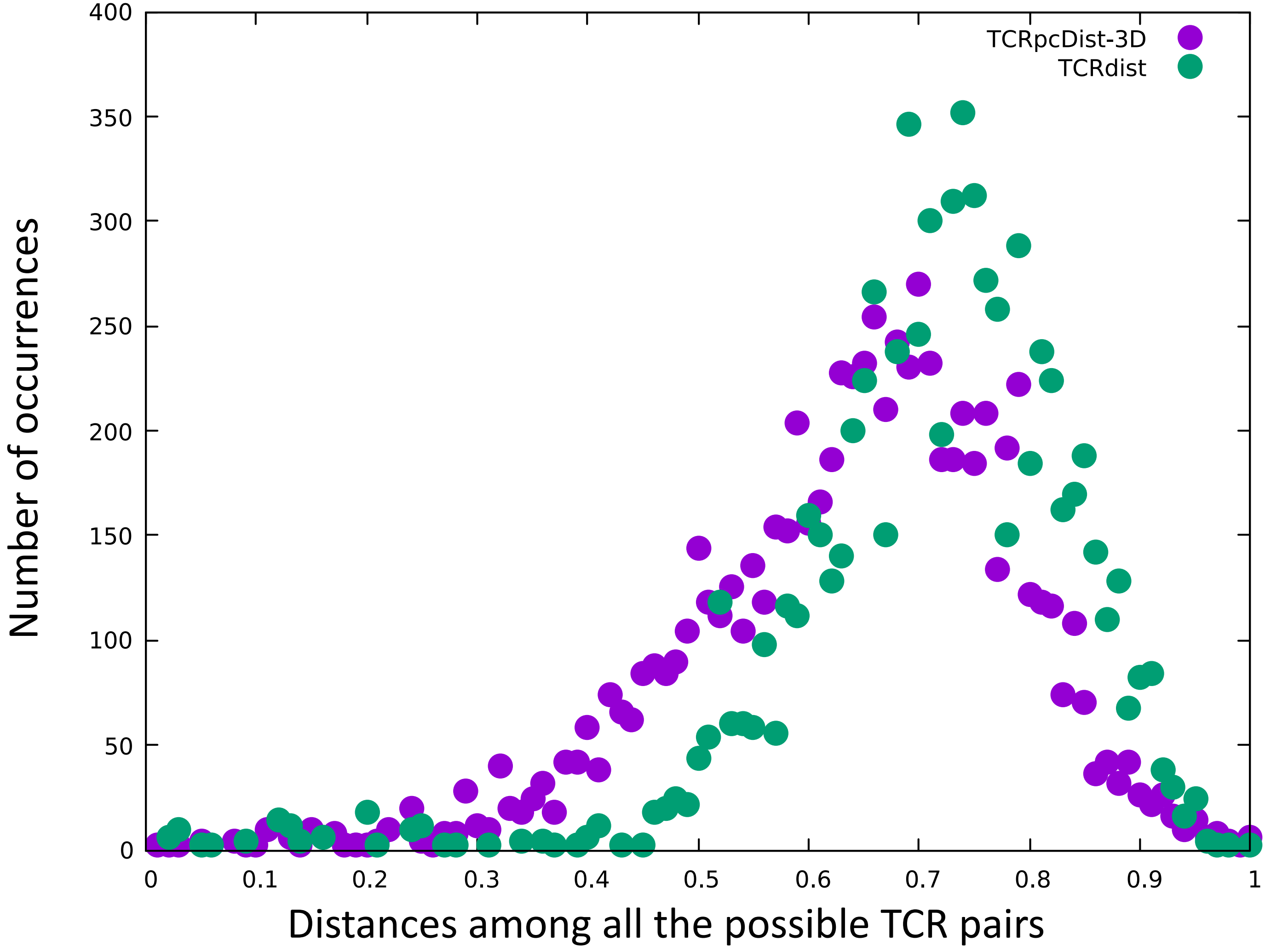


Higher the distances, the most different the TCRs

### PDB set of 84 TCRs - comparison between TCRpcDist-3D and TCRdist3

| | no threshold | | Threshold 0.15 | | | | Threshold 0.1 | | | |
|---|---|---|---|---|---|---|---|---|---|---|
| | TCRpcDist | TCRdist3 | TCRpcDist | | TCRdist3 | | TCRpcDist | | TCRdist3 | |
| | Success % | Success % | Success % | N. TCRs | Success % | N. TCRs | Success % | N. TCRs | Success % | N. TCRs |
| rank1 | **40.47** | 40.47 | 61.7 | **29** | 63.41 | 26 | 60.45 | **26** | 59.45 | 22 |
| rank2 | 42.85 | 45.23 | 65.95 | **31** | 68.29 | 28 | 65.11 | **28** | 64.86 | 24 |
| rank5 | **51.19** | 48.8 | 70.21 | **33** | 70.73 | 29 | 69.76 | **30** | 67.56 | 25 |
| rank10 | **54.76** | 54.76 | 72.34 | **34** | 73.17 | 30 | 69.76 | **30** | 70.27 | 26 |
| TCRs clustered | 84 | | 47 | | 41 | | 43 | | 37 | |
